# RIPK3-RNASE1 axis as a potential therapeutic and clinical monitoring target in VEXAS syndrome

**DOI:** 10.64898/2026.02.04.703672

**Authors:** Kana Higashitani, Tatsuma Ban, Yohei Kirino, Go R Sato, Soichiro Adachi, Yuki Iizuka, Ayaka Maeda, Osamu Ohara, Hideaki Nakajima, Tomohiko Tamura

**Author notes:** K.H. and T.B. contributed equally as first authors. Y.K. and T.B. contributed equally.

## Abstract

Vacuole, E1 enzyme, X-linked, autoinflammatory, somatic (VEXAS) syndrome (VS) is a severe autoinflammatory disease driven by somatic mutations in ubiquitin-like modifier activating enzyme 1 (*UBA1*), for which disease activity biomarkers and therapeutic targets are needed. We conducted longitudinal deep phenotyping of patients with VS and found that ribonuclease 1 (*RNASE1*) expression is most strongly correlated with the disease activity score measured by the VEXAS Current Activity Form (VEXASCAF). Single-cell RNA-sequencing showed that *RNASE1* was upregulated in VS monocytes. We generated human monocytic cell lines harboring *UBA1* mutations (p.Met41Val, p.Met41Leu, and p.Met41Thr). These cells exhibited varying degrees of impaired ubiquitination and subsequent pathological features such as the unfolded protein response, increased pro-inflammatory cytokine production, cell death, and *RNASE1* expression, mirroring the genotype–phenotype associations observed in patients with VS. Multi-omics analyses revealed that genotypes linked to greater clinical severity were enriched in pro-inflammatory, interferon, and necroptosis signatures. Notably, functional interrogation demonstrated that inhibition of receptor-interacting protein kinase 3 (RIPK3), a key regulator of necroptosis, markedly suppressed all these pathological features, including the elevated *RNASE1* expression. These results identify the RIPK3–RNASE1 axis in VS, highlighting RNASE1 as a potential biomarker and RIPK3 as an attractive therapeutic target.

## Introduction

VEXAS syndrome (an acronym for vacuoles, E1 enzyme, X-linked, autoinflammatory, somatic syndrome, abbreviated to VS) is an autoinflammatory disease that affects older males and is attributed to acquired somatic mutations in the ubiquitin-like modifier activating enzyme 1 gene (*UBA1*) located on the X chromosome.^1^ The E1 enzyme UBA1, particularly the cytosolic UBA1b isoform, is essential for initiating ubiquitination, which modulates various cellular processes, including cell survival. Hypomorphic *UBA1* mutations are considered to cause autoinflammatory symptoms associated with the activation of innate immunity.^2^ Previous reports, including our prospective registry study that established the disease activity measure VEXAS Current Activity Form (VEXASCAF), demonstrated that VS is characterized by a low remission rate and a high frequency of complications, mainly owing to severe inflammatory relapses following treatment tapering.^3–7^

The most common mutation hotspots are the methionine (Met) 41 site of *UBA1* exon 3, including p.Met41Val (c.121A>G), p.Met41Thr (c.122T>C), and p.Met41Leu (c.121A>C),^8,9^ hereafter referred to as M41V, M41T, and M41L, respectively. These three mutations account for over 90% of all reported patients with VS identified to date.^10^ Among these, M41V is associated with the poorest prognosis.^11,12^ Conversely, M41L has been linked to a milder clinical course.^13^ In addition, low variant allele frequency (VAF) of M41L has been reported in individuals without particular symptoms and in females.^14^ These findings suggest that the genotype of the *UBA1* mutation affects the phenotype in VS.

Glucocorticoids, well-known to induce various side effects, including opportunistic infections, serve as the initial therapy for managing disease activity in VS. Although newer drugs, such as Janus kinase (JAK) and IL-6 inhibitors, are also effective,^15–17^ glucocorticoid-sparing agents are currently unavailable. Emerging chemical compounds, such as UBA1/6 inhibitors and the gold-containing drug auranofin, show promise but remain experimental.^18,19,20^ Moreover, therapeutic strategies based on the type of *UBA1* mutations have been lacking. Accordingly, novel therapies are urgently needed to improve the prognosis and quality of life of patients with VS, which can be achieved through a more comprehensive understanding of the disease mechanism and pathogenesis.

In the current study, we performed deep phenotyping (DP) by longitudinally analyzing gene expression data linked to clinical information. We investigated the molecular mechanism through which *UBA1* mutations induce cellular abnormalities using newly established VS model cell lines. Herein, ribonuclease 1 (RNASE1) and receptor-interacting protein kinase 3 (RIPK3) emerged as key molecules that could serve as a potential biomarker and an attractive therapeutic target, respectively.

## Results

### Clinical features of registered participants

We enrolled 13 patients with VS carrying pathogenic *UBA1* mutations, 24 patients with *UBA1* mutation-negative VS-like autoinflammatory diseases (VLAD), and 10 healthy controls (HC) (Figure 1A). Table 1 and Supplemental Table 1 summarize the distribution of age, sex, symptoms associated with VS at study enrollment, and *UBA1* genotypes. Consistent with previous reports, most patients with myelodysplastic syndromes (MDS) in the VS group had a low Revised International Prognostic Scoring System (IPSS-R) score (Supplemental Table 2).^22^ Similar to patients with VS, those with VLAD were typically older males presenting with systemic inflammatory symptoms and a severe clinical course, often complicated by MDS. We also included 37 sex-matched patients with BD as a disease control.

**Figure 1.**
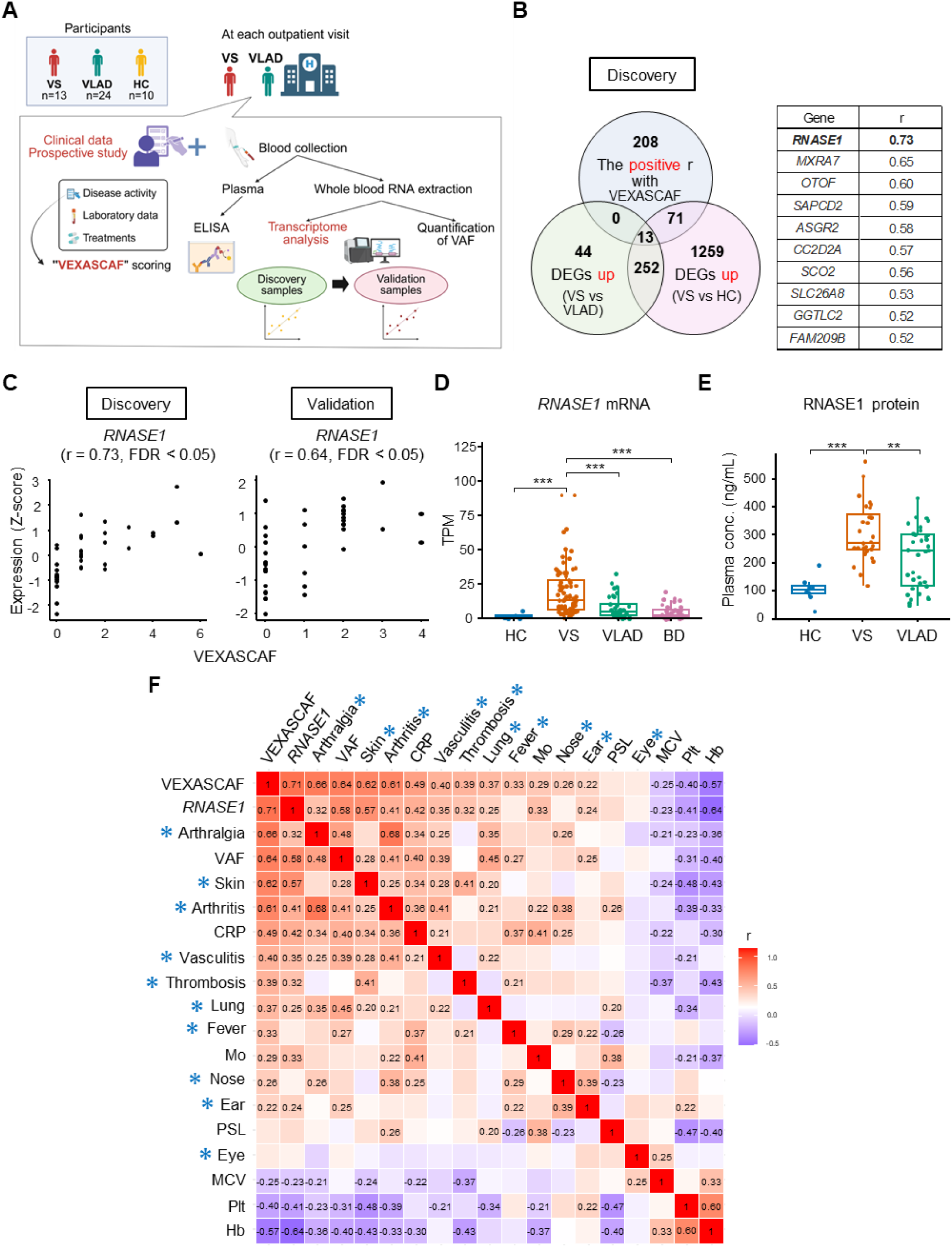
Deep phenotyping identifies RNASE1 as associated with disease activity in VS. **A:** Overview of the DP study design. Peripheral blood samples were collected from patients with VS (n = 13), VLAD (n = 24), and HC (n = 10) at each outpatient visit for RNA-seq, *UBA1* mutation analysis, VAF analysis, and plasma storage for ELISA. RNA-seq was performed separately during the discovery and validation phases. Clinical data were prospectively collected. DP, deep phenotype; VS, VEXAS syndrome; VLAD, VEXAS-like autoinflammatory disease; HC, healthy controls; VAF, variant allele frequency. **B:** Identification of *RNASE1* using DP. Genes positively correlated with VEXASCAF, DEGs between VS and VLAD, and VS and HC. Genes common to all comparisons are ranked by their correlation with VEXASCAF. DEGs were defined as FC > 2, FC < 1/2, and FDR < 0.05. VEXASCAF, VEXAS Current Activity Form; DEGs, differentially expressed genes. **C:** Scatter plots illustrating the correlation between *RNASE1* expression and VEXASCAF in both discovery and validation samples. **D:** Comparison of *RNASE1* expression levels among HC (n = 10), VS (n = 79), BD (n=49) and VLAD (n = 36) based on longitudinal RNA-seq analysis. Outliers (Vo) were defined as values meeting the criterion: Vo ≤ Q1 − 3 × IQR or Vo ≥ Q3 + 3 × IQR, where Q1 and Q3 are the first and third quartiles, and IQR is the interquartile range. Outliers were excluded from the analysis. BD, Behçet’s disease. **E:** Plasma RNASE1 protein levels measured by ELISA in HC (n = 8), VS (n = 28), and VLAD (n = 34) samples. **F:** Heatmap showing associations between VEXASCAF score, *RNASE1* gene, VAF of *UBA1* mutation, and various clinical parameters in patients with VS. The clinical symptoms assessed using the VEXASCAF scoring system are marked with asterisks. “Eye” includes ocular inflammation and periorbital edema; “Ear” and “Nose” indicate auricular and nasal chondritis, respectively. Values represent Spearman’s correlation coefficient (r) with *P* values < 0.05. *\*P* < 0.05*, **P* < 0.01*, ***P* < 0.001, ns: not significant. Box-and-whisker plots represent the median, IQR, and 1.5× IQR. Statistical comparisons were performed using the two-sided Mann–Whitney U test, and correlations were evaluated using two-sided Spearman’s rank correlation.

**Table 1.** Clinical characteristics of participants registered in deep phenotyping. Demographic, genetic, clinical, laboratory, and treatment data at study entry are summarized for healthy controls (HC), patients with Behçet’s disease (BD), VEXAS-like autoinflammatory disease (VLAD), and VEXAS syndrome (VS). Data are presented as numbers (%) for categorical variables and median (range or interquartile range, IQR) for continuous variables. *P* values were calculated using the Mann–Whitney U test for continuous variables and Fisher’s exact test for categorical variables. ‘Splice’ refers to the c.118-1G>C mutation. MCV, mean corpuscular volume; NA, not available; DMARDs, disease-modifying antirheumatic drugs. Synthetic DMARDs: azathioprine, cyclosporine, tacrolimus, sulfasalazine, and methotrexate. Biologic DMARDs: tocilizumab and infliximab.

|  | HC | BD | VLAD | VS | *P*-value<br>(VLAD and VS) |
| --- | --- | --- | --- | --- | --- |
| Demographic characteristics of participants |  |  |  |  |  |
| Participants, n | 10 | 37 | 24 | 13 |  |
| Deep phenotyping sample, n | 10 | 49 | 36 | 79 |  |
| Male sex, n (%) | 6 (60) | 37 (100) | 24 (100) | 13 (100) | 1 |
| Median age at study entry (range) | 35 (30–57) | 48 (19–75) | 74.5 (43–87) | 73 (55–88) | 1 |
| Genetic characteristics |  |  |  |  |  |
| p.Met41Val (c.121A > G), n (%) | NA | NA | 0 (0) | 2 (15.4) |  |
| p.Met41Thr (c.122T > C), n (%) | NA | NA | 0 (0) | 4 (30.8) |  |
| p.Met41Leu (c.121A > C), n (%) | NA | NA | 0 (0) | 5 (38.5) |  |
| Splice, n (%) | NA | NA | 0 (0) | 1 (7.7) |  |
| p.Ala478Ser, n (%) | NA | NA | 0 (0) | 1 (7.7) |  |
| Diagnosis |  |  |  |  |  |
| Myelodysplastic syndrome, n (%) | 0 (0) | 0 (0) | 8 (33.3) | 5 (38.5) | 1 |
| Relapsing polychondritis, n (%) | 0 (0) | 0 (0) | 0 (0) | 7 (53.8) | < 0.05 |
| Sweet's syndrome, n (%) | 0 (0) | 0 (0) | 0 (0) | 3 (23.1) | < 0.05 |
| Giant-cell arteritis, n (%) | 0 (0) | 0 (0) | 2 (8.3) | 1 (7.7) | 1 |
| Deaths, n (%) | 0 (0) | 0 (0) | 5 (20.8) | 1 (7.7) | 0.395 |
| Clinical features at study entry |  |  |  |  |  |
| Fever, n (%) | 0 (0) | NA | 10 (41.7) | 3 (23.1) | 0.305 |
| Ear chondritis, n (%) | 0 (0) | 0 (0) | 0 (0) | 2 (15.4) | 0.117 |
| Nose chondritis, n (%) | 0 (0) | 0 (0) | 0 (0) | 2 (15.4) | 0.117 |
| Lung symptoms, n (%) | 0 (0) | 0 (0) | 5 (20.8) | 0 (0) | 0.140 |
| Arthralgia, n (%) | 0 (0) | 4 (10.8) | 9 (37.5) | 5 (38.5) | 1 |
| Arthritis, n (%) | 0 (0) | 0 (0) | 2 (8.3) | 3 (23.1) | 0.321 |
| Thrombosis, n (%) | 0 (0) | NA | 1 (4.2) | 2 (15.4) | 0.278 |
| Dermatosis, n (%) | 0 (0) | 16 (43.2) | 7 (29.2) | 1 (7.7) | 0.216 |
| Eye inflammation, n (%) | 0 (0) | 1 (2.7) | 4 (16.7) | 0 (0) | 0.276 |
| Periorbital edema, n (%) | 0 (0) | NA | 1 (4.2) | 0 (0) | 1 |
| Vasculitis, n (%) | 0 (0) | 0 (0) | 1 (4.2) | 2 (15.4) | 0.278 |
| Median VEXASCAF (range) | 0 (0) | NA | 1.5 (0-5) | 1 (0-6) | 0.637 |
| Laboratory data at study entry |  |  |  |  |  |
| Hemoglobin, g/dL (IQR) | NA | 14.4 (13.9–15) | 10.4 (9.1–11.5) | 9.4 (8.7–12.1) | 0.820 |
| MCV, fL (IQR) | NA | 91.1 (89.5–94.7) | 94.8 (89.6–107) | 106 (101–109) | < 0.05 |
| Leukocytes, ×10<sup>9</sup>/L (IQR) | NA | 6.5 (4.9–6.8) | 4.4 (3.0–6.9) | 4.5 (3.0–6.1) | 0.772 |
| Neutrophils, ×10<sup>9</sup>/L (IQR) | NA | 3.3 (2.5–4.1) | 2.8 (1.6–5.3) | 3.4 (2.0–3.9) | 0.790 |
| Monocytes, ×10<sup>9</sup>/L (IQR) | NA | 0.53 (0.39–0.67) | 0.56 (0.33–0.83) | 0.27 (0.18–0.35) | < 0.05 |
| Platelets, ×10<sup>3</sup>/μL (IQR) | NA | 259 (214–289) | 217 (122–294) | 127 (82–206) | 0.227 |
| C-reactive protein, mg/L (IQR) | NA | 1.0 (0.2–2.8) | 23 (2.8–79.3) | 3.0 (0.2–35) | 0.157 |
| Treatment at study entry |  |  |  |  |  |
| Glucocorticoids, n (%) | 0 (0) | 5 (13.5) | 14 (58.3) | 9 (69.2) | 0.725 |
| Median prednisone dose, mg/day (range) | 0 (0) | 0 (0) | 15 (0–40) | 9 (0–40) | 0.789 |
| Synthetic DMARDs, n (%) | 0 (0) | 13 (35.1) | 3 (12.5) | 0 (0) | 0.538 |
| Biologic DMARDs, n (%) | 0 (0) | 26 (70.3) | 5 (20.8) | 4 (30.8) | 0.691 |

### Identification of molecules most correlated with VS disease activity through DP

We performed DP to identify molecules associated with the VS disease activity index (DAI). Longitudinal bulk RNA sequencing (RNA-seq) analysis (n = 79 samples) was performed on two independent occasions, with the first group in time defined as the discovery samples (38 samples) and the latter group as the validation samples (41 samples; Supplemental Table 3, Supplemental Figure 1).

We extracted the differentially expressed genes (DEGs) from the VS, VLAD, and HC discovery whole blood RNA samples. Thereafter, we screened for genes upregulated in VS that were positively correlated with DAI using VEXASCAF. Among these genes, *RNASE1* exhibited the highest correlation coefficient (Figure 1B; Supplemental Figure 2A). A similar analysis of the validation samples confirmed that *RNASE1* was the top-ranked gene (Figure 1C; Supplemental Figure 2A-B). The *RNASE1* mRNA level was significantly elevated in patients with VS compared with patients with VLAD, HC, or BD (Figure 1D). The level of RNASE1 in plasma, as measured using enzyme-linked immunosorbent assay (ELISA), showed a similar trend (Figure 1E). In addition to the factors assessed in VEXASCAF, *RNASE1* expression correlated with other clinical symptoms (Figure 1F). Specifically, *RNASE1* expression positively correlated with the VAF of *UBA1* mutation and negatively with hemoglobin, indicating that clonal and ineffective hematopoiesis were enhanced in patients with high VS disease activity. These data suggested that RNASE1 expression was associated with VS disease activity.

### Further characterization of RNASE1 expression in VS

Next, we analyzed serum samples previously collected during treatment.^23^ The results revealed a decrease in RNASE1 concentration accompanied by improvement in symptoms following an increased dose of PSL (Figure 2A; Supplemental Table 4). Proteomic analysis of plasma from separate patients with VS (Supplemental Table 5) also revealed the presence of elevated RNASE1 levels in VS (Supplemental Figure 3).

**Figure 2.**
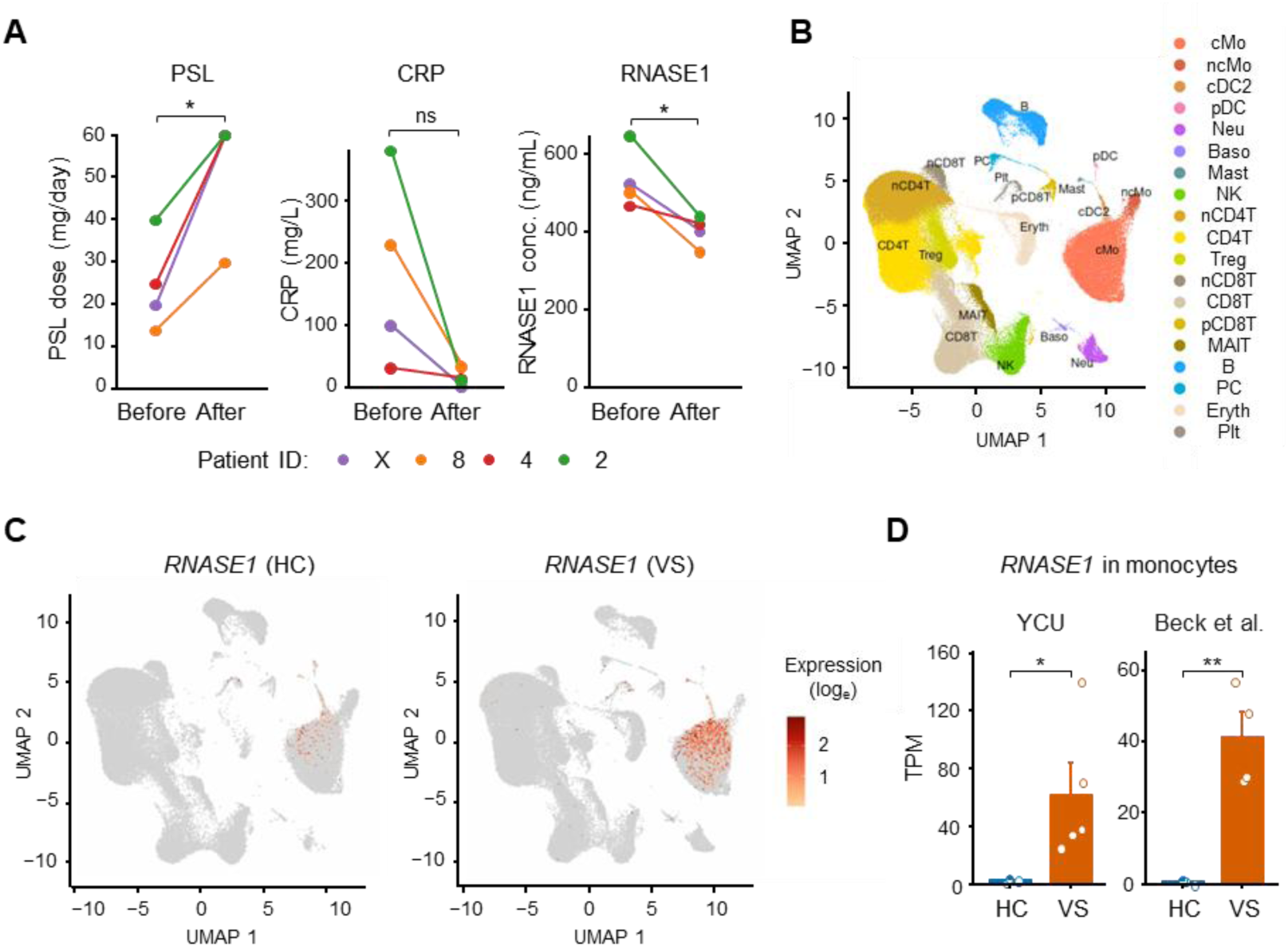
Further characterization of RNASE1 expression in patients with VS. **A:** Serum RNASE1 levels before and after prednisolone (PSL) dose escalation in the patients with VEXAS syndrome (VS). Changes in PSL dosage, C-reactive protein (CRP) levels, and serum RNASE1 concentrations are shown for four patients with VS who underwent PSL dose increases owing to worsening disease activity. RNASE1 levels significantly decrease following PSL escalation. **B:** Uniform Manifold Approximation and Projection (UMAP) plot of 253,296 cells in peripheral blood mononuclear cells (PBMCs) from Yokohama City University (YCU) (HC; n = 1, VS; n = 2) and Mizumaki et al (HC; n = 5, VS; n = 9). Cell type annotations for each cluster are indicated. Mo, monocyte; cMo, classical Mo; ncMo, non-classical Mo; DC, dendritic cells; cDC, conventional DC; pDC, plasmacytoid DC; Neu, neutrophil; Baso, basophil; NK, natural killer; n, naïve; Treg, regulatory T; p, proliferating; MAIT, mucosal-associated invariant T; PC, plasma cell; Eryth, erythrocyte; Plt, platelet. **C:** UMAP plots of PBMCs derived from HC (n = 6) and VS patients (n = 11), colored according to *RNASE1* expression levels in each dataset. The total cell numbers were downsampled to 50,000 cells per group. **D:** Comparison of *RNASE1* expression levels in peripheral blood monocytes between HC and VS groups using bulk RNA sequencing. Left: Data from YCU (HC: n = 3, VS: n = 5); Right: Independent dataset from Beck, et al (HC: n = 3, VS: n = 4). *\*P* < 0.05*, **P* < 0.01*, ***P* < 0.001, ns: not significant. Error bars indicate the standard error of the mean (**D**). Statistical significance was assessed using a two-sided paired *t*-test (**A**) and a two-sided t-test (**D**).

To understand which cell types express *RNASE1*, we performed single-cell RNA sequencing (scRNA-seq) of peripheral blood mononuclear cells (PBMCs) from HC or patients with VS and the data were integrated with those by Mizumaki et al.^24^ We found that classical monocytes from patients with VS highly expressed *RNASE1* (Figure 2B, C, and Supplemental Figure 4A). In addition, bulk RNA-seq analysis of monocytes from our patients and a publicly available dataset by Beck et al.^1^ confirmed the elevated expression of *RNASE1* in monocytes from patients with VS (Figure 2D). Similarly, scRNA-seq data on bone marrow mononuclear cells from another publicly available dataset^25^ revealed high *RNASE1* expression in VS monocytes (Supplemental Figure 4B and 5). Interestingly, *RNASE1* expression was also detected in hematopoietic stem/progenitor cells, which may warrant future investigation. These findings from clinical samples indicated that high *RNASE1* expression in monocytes is associated with increased disease activity in VS.

To examine the relationship between *RNASE1* expression and clonal hematopoiesis of indeterminate potential (CHIP), we analyzed a publicly available RNA-seq dataset of older adult monocytes.^26^ We found that *RNASE1* expression was not associated with CHIP (Supplemental Figure 6). We also reviewed DEG results from publicly available whole-blood RNA-seq datasets of healthy individuals,^27,28^ which indicated that neither aging nor sex was associated with increased *RNASE1* expression.

### Differences in *RNASE1* expression based on genotype and VAF

DP RNA-seq data, including five *UBA1* mutations, were used for principal component analysis (PCA) to determine the correlation between the genotype and phenotype. A distinct gene expression pattern was associated with the *UBA1* genotype (Figure 3A), with the highest *RNASE1* expression associated with M41L, while the lowest was linked to M41V (Figure 3B). Consistent with the strong association between VEXASCAF and *RNASE1*, the distribution of VEXASCAF by genotype closely mirrored that of *RNASE1* (Figure 3C). The peripheral blood *UBA1* mutant VAF in DP samples exhibited distinct genotypic differences (Figure 3D). The higher VAF in the M41L mutation was consistent with our previous findings.^3,21^

**Figure 3.**
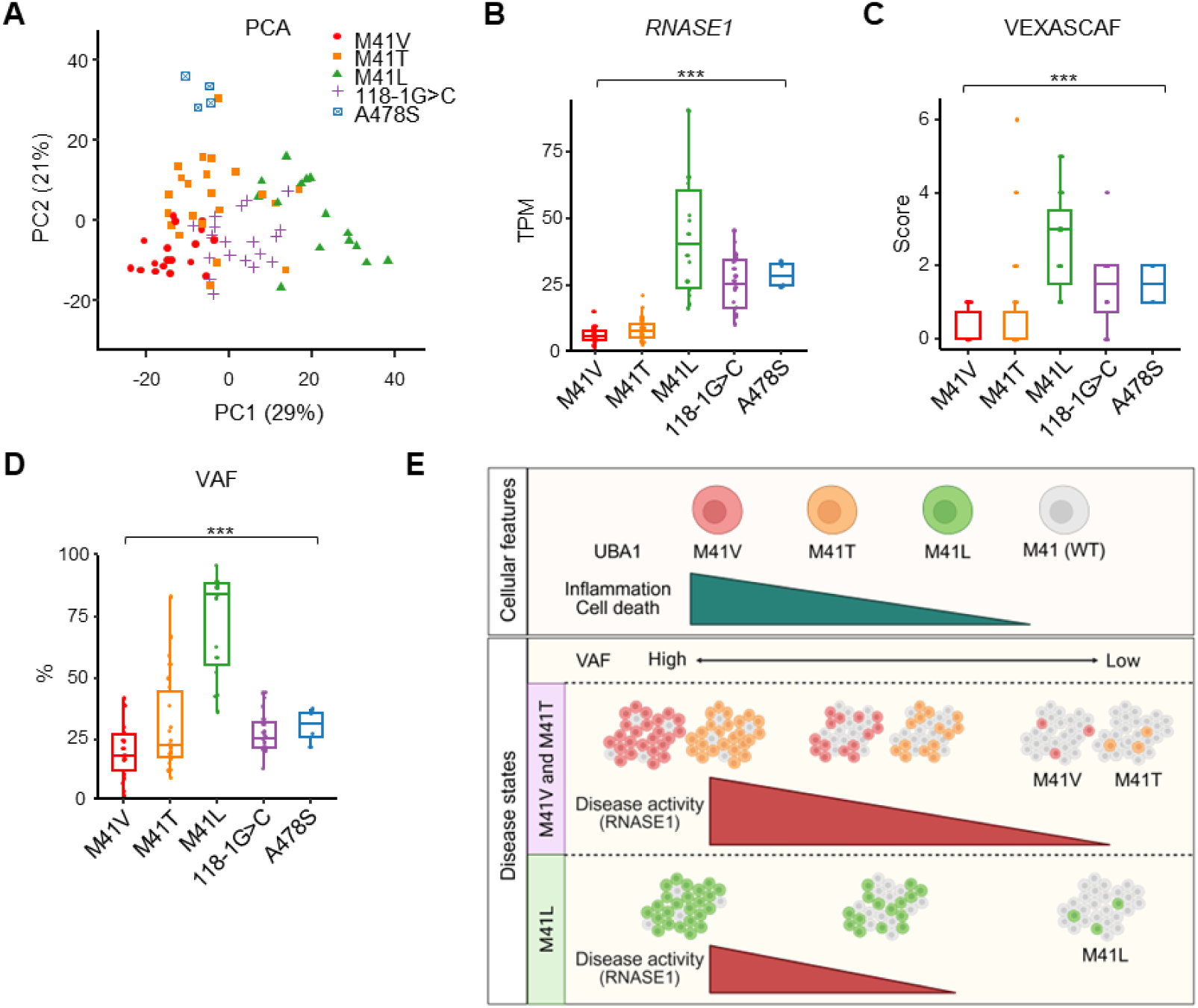
*UBA1* genotype-dependent differences in patients with VS. **A:** Principal component analysis (PCA) of RNA sequencing data from patients with VEXAS syndrome (VS) who participated in deep phenotyping (DP) (n = 79), stratified by *UBA1* genotype. **B:** *RNASE1* expression levels in transcripts per million (TPM) in patients with VS who participated in DP stratified by *UBA1* genotype. Outliers (Vo) were defined as values meeting the criterion: Vo ≤ Q1 − 3 × IQR or Vo ≥ Q3 + 3 × IQR. A graph that excluded these outliers was generated. **C:** VEXASCAF score of patients with VS stratified by *UBA1* genotype. **D:** Variant allele frequency (VAF) (%) of patients with VS stratified by *UBA1* genotype. **E:** Schematic overview illustrating genotype-phenotype correlation and disease progression in VS. (Top) Different *UBA1* genotype clones (M41V, M41T, M41L, and wild-type [WT]) are associated with varying degrees of inflammation and cell death. (Bottom) Individuals with low *UBA1* mutant VAF and the M41L variant tend to exhibit low pathogenicity, whereas the M41V and M41T variants are associated with VS development. In contrast, moderate to high VAF is associated with the high pathogenicity of M41V and M41T, whereas M41L may lead to VS symptoms at moderate VAF. *P* values were determined using the two-sided Kruskal-Wallis test, followed by Dunn’s post-hoc test for multiple group comparisons (**B**-**D**). Box-and-whisker plots display the median, interquartile range (IQR), and 1.5× IQR.

The high disease activity in patients carrying M41L was slightly unexpected. Taken together with the genotype-dependent features of VS cell lines described in the following sections, we suggest a relationship between cellular abnormalities, VAF, and disease activity in VS (Figure 3E; see Discussion).

### *UBA1* genotype-dependent differences in VS cell lines

The above-described results prompted us to investigate the mechanisms underlying elevated *RNASE1* expression in VS monocytes. Therefore, human monocytic VS cell lines were generated. Specifically, we introduced three common *UBA1* mutations associated with VS, namely M41V, M41T, and M41L, into THP-1 cells, a widely used monocytic cell line with an XY sex chromosome (Figure 4A; Supplemental Figure 7).^29,30^ Because these *UBA1* mutations lead to the loss of the UBA1b isoform, resulting in cell death, we used the Tet-On/Off system to maintain the exogenous expression of UBA1b by Tet-On to obtain mutant clones. The VS state was induced by Tet-Off (without doxycycline [Dox]), leading to the depletion of exogenous UBA1b within 12 days (Supplemental Figure 8A), with UBA1a and UBA1c from the endogenous mutant allele remaining (Figure 4B). The expression of another E1 enzyme, UBA6, remained unchanged regardless of the Tet-On or Tet-Off conditions. Notably, although UBA1b levels in Tet-On VS cell lines were higher than endogenous UBA1b, RNA-seq data on M41M cells (M41-unmutated cells harboring Tet-On inducible *UBA1b*) with or without Dox detected almost negligible impact of UBA1b overexpression (Supplemental Figure 8B).

**Figure 4.**
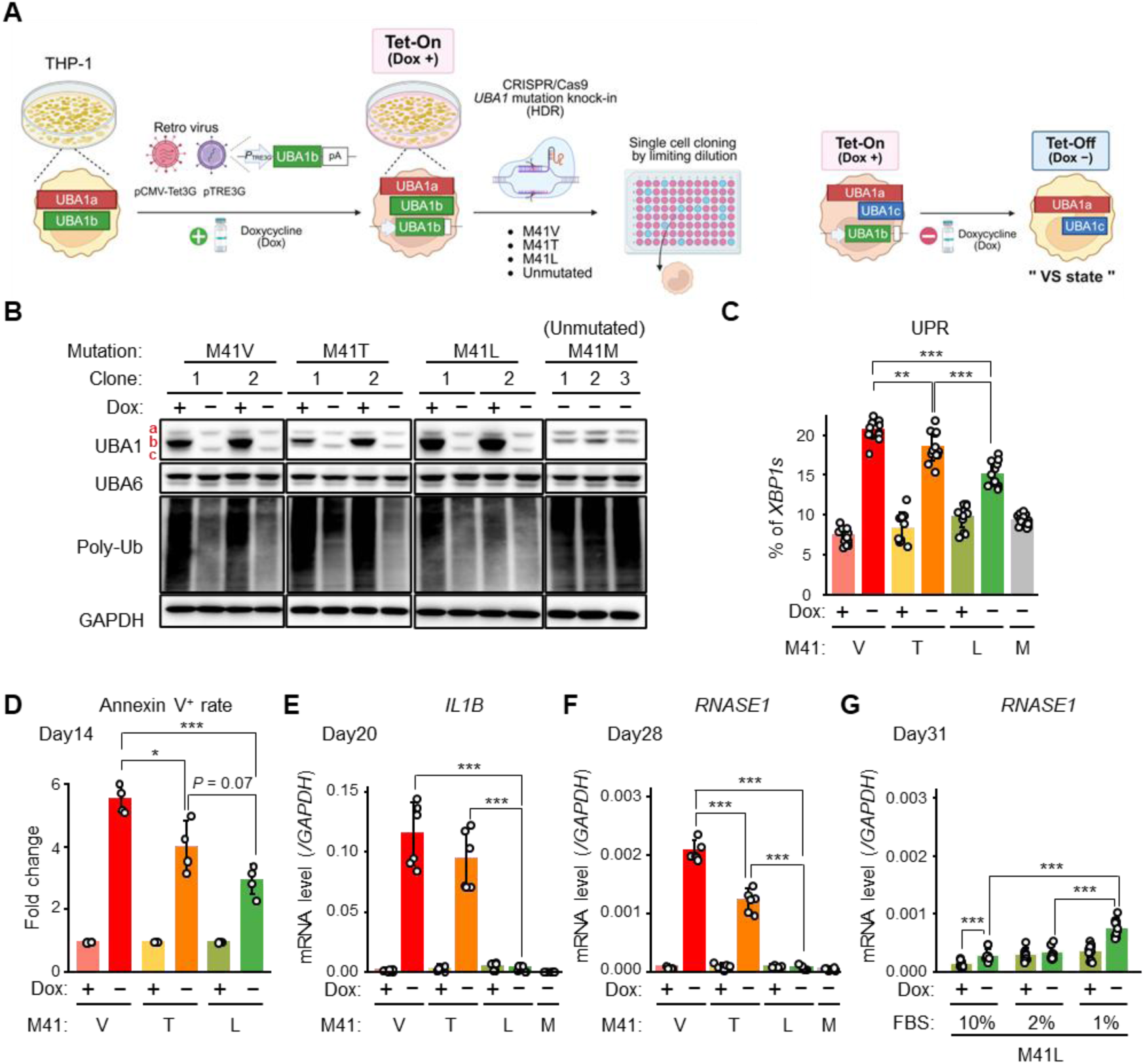
VS cell lines recapitulate key characteristics of primary VS cells. **A:** Schematic diagram illustrating the VEXAS syndrome (VS) cell line generation method. In addition to *UBA1* mutations associated with VS (M41V, M41T, and M41L), we generated a control cell line with a silent mutation near the UBA1 Met41 position, which is referred to as “Unmutated.” **B:** Immunoblot analysis of UBA1, UBA6, and polyubiquitin proteins. GAPDH was used as a loading control. Experiments were performed on day 19 after the switch to Tet-On (Dox +, 20 ng/mL) or Tet-Off (Dox −) conditions. Dox, doxycycline. **C:** Results of XBP1 splicing measurements show an increased percentage of spliced XBP1 with genotype-specific differences. Cell lines were used for experiments on day 18 after switching. **D:** Annexin V/propidium iodide assays showed an increase in Annexin V-positive dead cells exhibiting membrane damage under Tet-Off conditions, with differences observed among genotypes. Assays were conducted 14 days after switching. **E:** mRNA expression levels of *IL1B* under Tet-On/Off conditions, measured on day 20. **F:** mRNA expression levels of *RNASE1* under Tet-On/Off conditions, measured on day 28. **G:** *RNASE1* mRNA expression in M41L VS cells cultured under serum starvation, measured on day 31 post-switching. Cell lines were cultured for six days in media containing varying fetal bovine serum (FBS) concentrations, starting from day 25 post-switching. Unmutated cells are labeled as “M41: M”. Data are representative of more than three independent repeat experiments (**B**). Data were pooled from two to four independent experiments using different clones (**C-G**). *\*P* < 0.05*, **P* < 0.01*, ***P* < 0.001, ns: not significant. Error bars indicate the standard deviation (SD). *P* values were determined using a two-sided *t*-test (**C-G**).

Clones carrying the VS mutations showed decreased viability and reduced proliferation after culturing under Tet-Off conditions for 12–14 days (Supplemental Figure 8C). Furthermore, immunoblotting analysis revealed impaired ubiquitination of these cells under Tet-Off conditions (Figure 4B). Cell morphology using Wright–Giemsa staining revealed an increase in the number of cytoplasmic vacuoles (Supplemental Figure 8D). These findings demonstrated that the phenotypic characteristics of our VS cell lines aligned with previously reported data from primary VS cells.^1,11^ Consistent with previous reports,^18,20^ our VS cell lines also exhibited high sensitivity to low concentrations of the UBA1 inhibitor, TAK-243 (Supplemental Figure 8E-F). It should be noted that the extent of reduced viability in M41L cells under Tet-Off conditions was more gradual than that in the other genotypes (Supplemental Figure 8C).

Because cell death observed in monocytes of patients with VS is linked to the activation of the unfolded protein response (UPR),^1,31^ we examined the percentage of the spliced form of X-box binding protein 1 (*XBP1*) mRNA (*XBP1s*), a well-known indicator of UPR activation. Under Tet-Off conditions, UPR activation was observed in all the mutants, demonstrating the following order of strength: M41V > M41T > M41L (Figure 4C). In support of this genotype-dependent difference, M41L cells exhibited a mild reduction in ubiquitination, which was more clearly observed when the cytoplasmic fraction was analyzed (Supplemental Figure 9A). In addition, the induction of Annexin V-positive cell death by Tet-Off was milder in M41L cells than in M41V and M41T cells (Figure 4D). Furthermore, the inflammatory gene *IL1B* was induced under Tet-Off conditions in M41V and M41T cells, whereas this induction was substantially lower in M41L cells (Figure 4E). Notably, *RNASE1* also showed a similar expression pattern (Figure 4F).

Temporal gene expression profiling revealed that *IL1B* expression in M41V and M41T cells was elevated as early as day 18, following Tet-Off induction, whereas *RNASE1* upregulation was observed after day 21, suggesting later involvement of RNASE1 in the inflammatory cascade (Supplemental Figure 9B-C). Consistent with the results shown in Figure 4F, M41L cells exhibited markedly lower *RNASE1* expression at the examined time points compared with other genotypes (Supplemental Figure 9C). Conversely, elevated *RNASE1* expression was observed in peripheral blood samples from patients with the M41L mutation (Figure 3B). Based on this discrepancy, we hypothesized that *RNASE1* expression is conditionally regulated *in vitro*. Indeed, long-term culture (31 days) and nutrient starvation under low fetal bovine serum conditions both significantly increased *RNASE1* expression, even in M41L cells (Figure 4G). Upon stimulation with nigericin, a signal 2 activator of the inflammasome, interleukin (IL)-1β and IL-18 secretion significantly increased in all genotypes (Supplemental Figure 9D-E). These results indicated that *UBA1* genotype-dependent differences in the phenotype and *RNASE1* expression may be initiated by the distinct extent of reduced ubiquitination. Notably, shRNA-mediated gene silencing (hereafter referred to as “knockdown”) revealed no RNASE1-mediated effect on cell death or *IL1B* expression (Supplemental Figure 10).

### Capturing early pathological changes of VS through omics analysis of VS cell lines

Considering the results of the temporal gene expression profiling and knockdown experiments, we examined early molecular changes occurring prior to the upregulation of *RNASE1* during VS induction. To obtain gene expression and chromatin accessibility profiles, we performed RNA-seq and an assay for transposase-accessible chromatin using sequencing (ATAC-seq) in VS cell lines on day 18 following Tet-Off conditions. PCA of the RNA-seq data revealed distinct transcriptional profiles between the Tet-Off and Tet-On conditions, with the degree of separation by PC1 ordered as follows: M41V > M41T > M41L (Figure 5A). Consistently, the numbers of DEGs from the RNA-seq data and differentially accessible regions (DARs) from the ATAC-seq data, comparing Tet-Off and Tet-On, were also ordered as M41V > M41T > M41L (Supplemental Figure 11A-B). Gene Set Enrichment Analysis (GSEA) using the MSigDB database^32^ revealed that M41V and M41T cells exhibited significant enrichment of genes related to interferon (IFN) and inflammatory responses in Tet-Off compared with Tet-On, consistent with previous data from patients with VS.^1^ In contrast, M41L cells demonstrated no significant gene set enrichment, indicating minimal transcriptional changes (Figure 5B).

**Figure 5.**
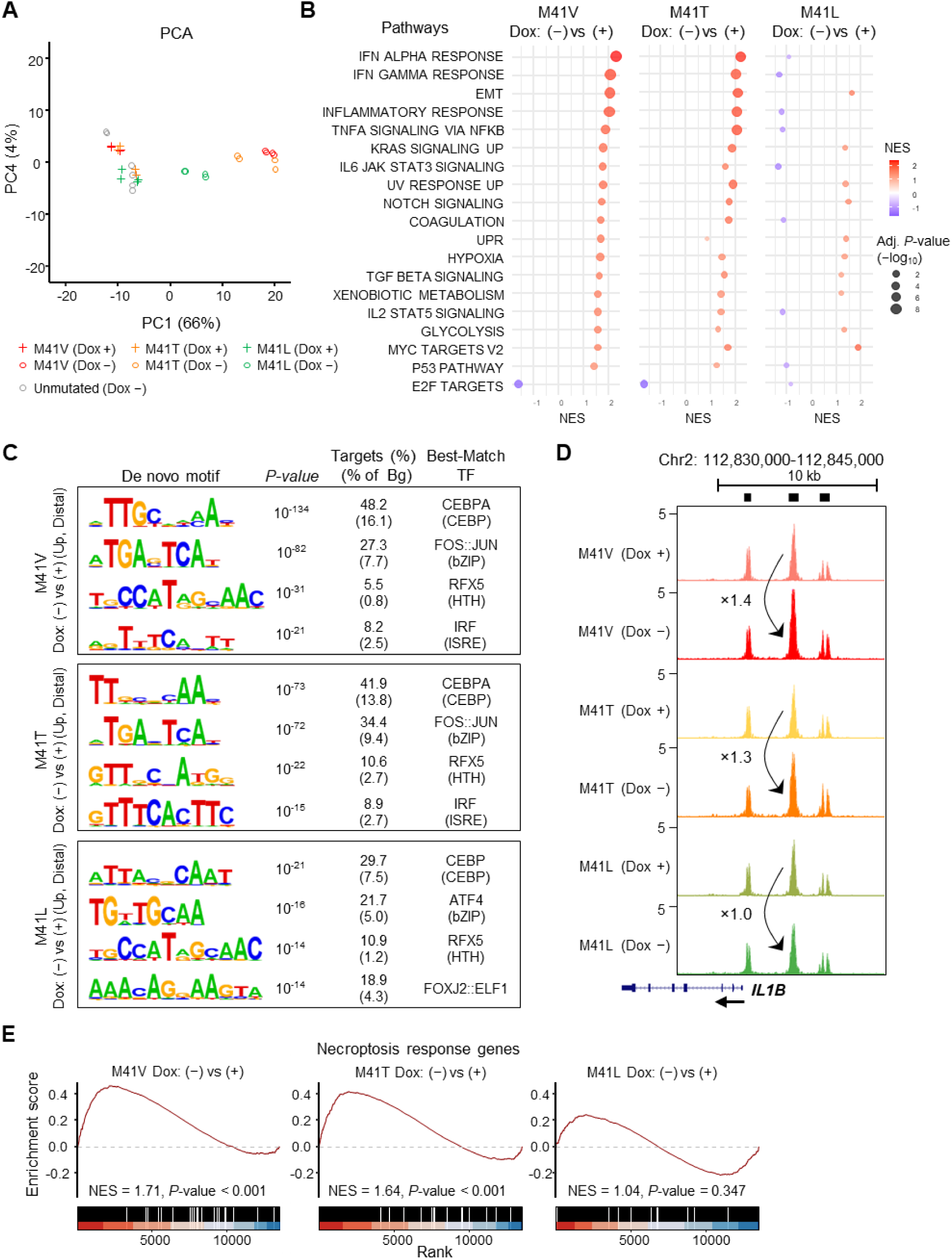
Multi-omics analysis of VS cell lines to reveal early pathogenic changes. **A:** Principal component analysis (PCA) of transcriptomic data from VEXAS syndrome (VS) and unmutated cell line samples, showing separation of both the genotype and the Tet-On (Dox +)/Off (Dox −) conditions primarily along principal component 1 (PC1). Two independent clones were used for each VS genotype, and three clones were used for the unmutated cell lines. Technical replicates were included for all the clones. **B:** Gene Set Enrichment Analysis (GSEA) comparing Tet-Off vs. Tet-On conditions in VS cell lines with different UBA1 genotypes. IFN, interferon; EMT, epithelial–mesenchymal transition; UPR, unfolded protein response. **C:** Integration of ATAC-sequencing and RNA sequencing data using Homer software. Differentially accessible regions (DARs) and differentially expressed genes (DEGs) were identified under Tet-Off vs. Tet-On conditions. A de novo motif analysis was conducted on peaks located at +2,000 or −2,000 base pairs relative to the transcription start site (TSS) of nearby genes, and regions exhibiting concordant upregulation of both DARs and DEGs were identified. Motif enrichment was quantified using binomial tests, and the results are reported as corresponding *P* values and motif frequency ratios. Bg, background. **D:** UCSC Genome Browser images of ATAC-seq data at the IL1B locus under Tet-On/Off conditions for each UBA1 genotype. Data were obtained from two independent clones per genotype, each analyzed in technical duplicates, and averaged for visualization according to genotype and condition. Black boxes above the ATAC-seq data tracks indicate merged peak calls across samples. **E:** GSEA of necroptosis response genes^34^ comparing Tet-Off vs. Tet-On conditions in VS cell lines with different UBA1 genotypes.

To identify the transcription factors involved in VS induction, we performed de novo DNA motif analysis of DARs. Approximately 130,000 ATAC-seq peaks were classified into proximal (less than 2,000 bp from the transcription start site [TSS]) and distal (2,000 bp or more from the TSS) peaks. Proximal peaks showing increased chromatin accessibility and upregulated nearest gene expression in the Tet-Off versus Tet-On conditions were enriched for a limited number of motifs. Conversely, distal peaks under the same comparison showed AP-1 (FOS::JUN) and IFN regulatory factor (IRF) motif enrichment in M41V and M41T cells but not in M41L cells (Figure 5C). These results suggested that, upon severe VS induction, IRF and AP-1 activate IFN and other inflammatory response pathways. As an example of an inflammation-related gene upregulated in the VS state, we confirmed the presence of a distal ATAC-seq peak for *IL1B*, which was upregulated under Tet-Off conditions (Figure 5D).

### RIPK3-dependent cell death in VS cell lines

Pronounced cell death was observed in both VS cell lines (Figure 4D, Supplemental Figure 8C) and in monocytes from patients with VS.^23,31^ The three most well-explored forms of programmed cell death are apoptosis, necroptosis, and pyroptosis. Apoptosis is a “silent,” inflammation-free form of cell death that is dependent on caspases, such as Caspase-3. Conversely, necroptosis, involving RIPK1, RIPK3, and MLKL, and pyroptosis, including Caspase-1 cleavage of gasdermin-D, lead to membrane rupture and inflammatory cell death.^33^ Among them, data on IL-1β and IL-18 secretion (Supplemental Figure 9D-E) suggested that pyroptosis was not executed in VS cell lines. Because the apoptosis-related gene sets in the MSigDB database were not enriched in VS cells (Supplemental Figure 11C), we performed GSEA on necroptosis response genes,^34^ revealing that Tet-Off M41V and M41T cells highly express these genes (Figure 5E). Moreover, we detected a sign of enhanced necroptosis, i.e., MLKL phosphorylation, in primary monocytes from the patient during disease flare, exhibiting a high VEXASCAF score (Supplemental Figure 12, Supplemental Table 6).

Our VS cell lines were treated with either a RIPK3 inhibitor, GSK’872, or a pan-caspase inhibitor Z-VAD-FMK (Z-VAD). GSK’872 inhibited necroptosis as evidenced by lower levels of phosphorylated MLKL and reduced percentages of Annexin V^+^, membrane-damaged cells (Figure 6A, lanes 5-6; Figure 6B).^35^ Conversely, Z-VAD strongly inhibited caspase activity and augmented necroptosis (Supplemental Figure 13) (Figure 6A, lane 4; Figure 6B). These results indicated that VS cells employ RIPK3-dependent necroptosis. This cell death pathway appears to be enhanced by inhibition of caspase activity. Correspondingly, Caspase-8 has been shown to suppress RIPK3-dependent necroptosis.^36–39^

**Figure 6.**
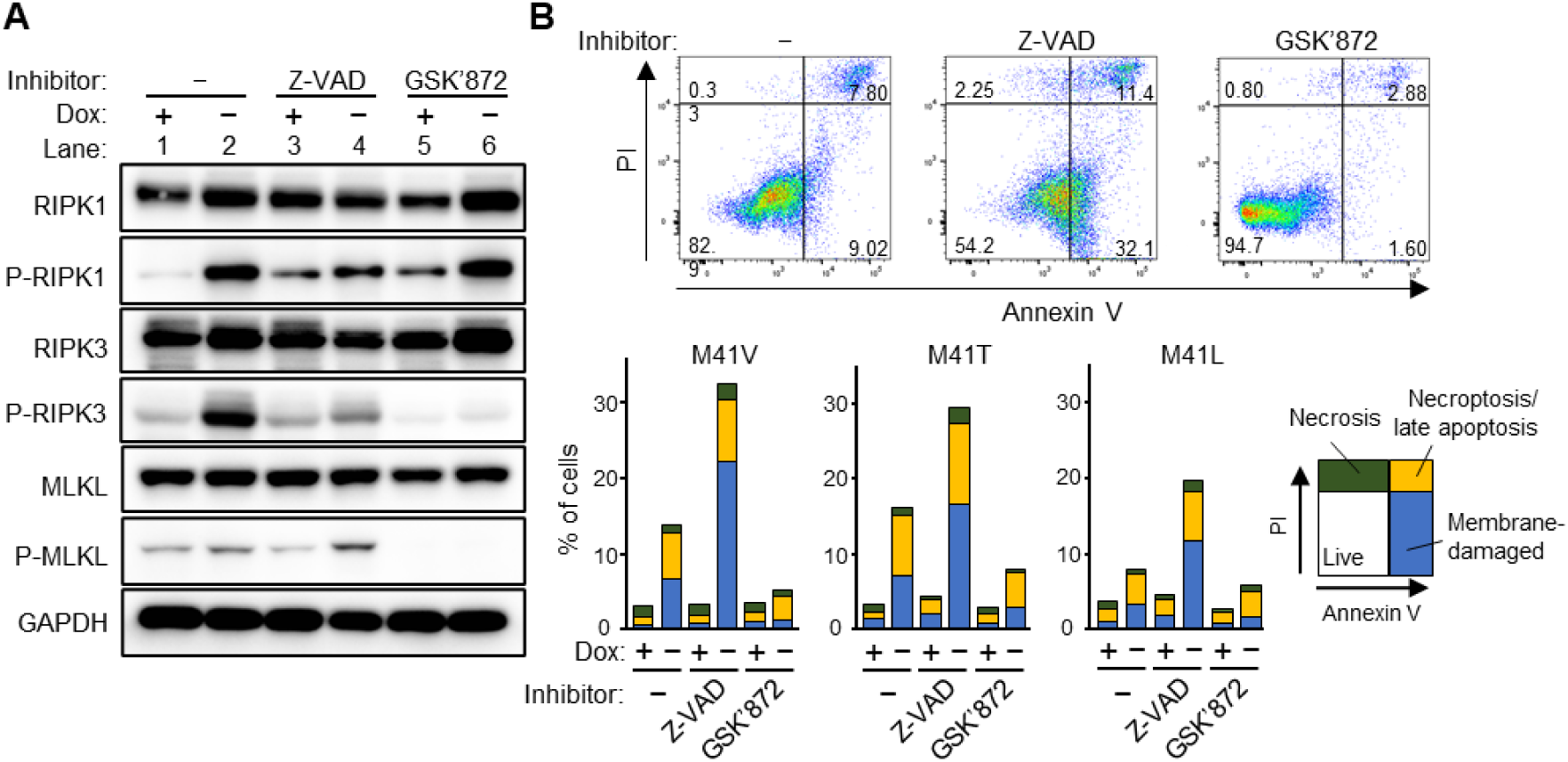
RIPK3-dependent cell death in VS cell lines. **A:** Immunoblot showing increased p-MLKL expression in the Tet-Off VEXAS syndrome (VS) cell line (M41V) following treatment with the pan-caspase inhibitor Z-VAD (10 µM), and decreased p-MLKL expression following treatment with the RIPK3 inhibitor GSK’872 (10 µM). The experiments were conducted on day 18 after switching to the Tet-Off condition. **B:** (Top) Representative Annexin V–propidium iodide (PI) flow cytometry plots of VS cell lines (M41V under Tet-Off conditions) cultured in medium containing DMSO, 10 µM Z-VAD, or 10 µM GSK’872. (Bottom) Percentages of cells showing membrane damage (phosphatidylserine exposure), late apoptosis/necroptosis, and necrosis in VS cell lines (M41V, M41T, and M41L under Tet-On/Off conditions) cultured in medium containing DMSO, Z-VAD, or GSK’872. Values represent the means of three independent experiments. The experiments were conducted 15 days after switching.

RNASE1 is reportedly released in response to extracellular RNA (eRNA), a component of damage-associated molecular patterns (DAMPs).^40^ Therefore, we investigated whether DAMPs released during necroptosis contribute to *RNASE1* expression by coculturing *UBA1*-unmutated (M41M) cells or alive VS cell lines with VS cell lines undergoing cell death. However, dying VS cells had no effect on the expression of *RNASE1* or *IL1B* in cocultured cells (Supplemental Figure 14). These results indicate that *RNASE1* upregulation occurs via a cell-intrinsic mechanism dependent on RIPK3, rather than in response to extracellular DAMPs.

### Key role of RIPK3 in various aspects of abnormalities in VS cell lines

To clarify the contribution of IRF, AP-1, and necroptosis to the pathological state of VS, we treated M41V cell lines with pharmacological inhibitors: anifrolumab (anti-IFNAR1 monoclonal antibody) for IRF, SP600125 (JNK inhibitor) for AP-1, and Nec-1s (RIPK1 inhibitor) or GSK’872 (RIPK3 inhibitor) for necroptosis. Targeted gene expression analysis revealed that IFN-stimulated genes (ISGs) *IFI27*, *MX1*, and *CXCL10* (encoding IP-10) were markedly suppressed by anifrolumab and GSK’872 (Figure 7A). In contrast, *IL1B* and *RNASE1* were significantly reduced by SP600125, Nec-1s, and GSK’872 (Figure 7B). Among these inhibitors, GSK’872 exhibited the broadest suppression of early VS pathological abnormalities.

**Figure 7.**
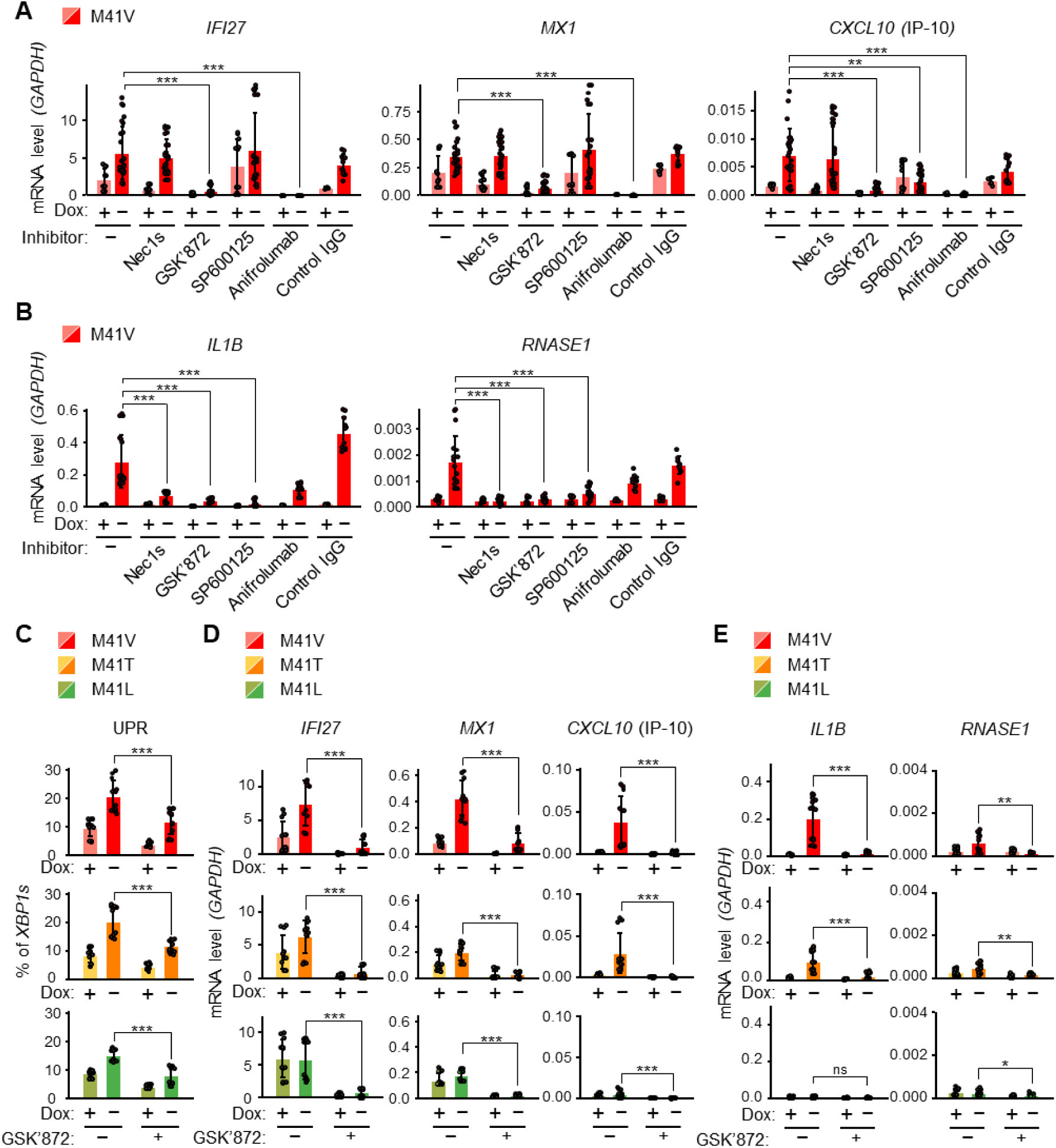
Involvement of RIPK3 in various abnormalities in VS cell lines. **A:** mRNA expression levels of *IFI27, MX1,* and *CXCL10* were measured after treatment with 30 µM Necrostatin-2 racemate (Nec-1s), 10 µM GSK’872, 10 µM SP600125, 10 µg/mL anifrolumab, or a 10 µg/mL human IgG1 isotype control. Experiments were conducted on M41V cells 18 days after switching. **B:** mRNA expression levels of *IL1B* and *RNASE1* were measured after treatment with the same inhibitors as in (**A**). Experiments were conducted on M41V cells 22 days after switching. **C:** The proportion of spliced XBP1 decreased upon treatment with GSK’872. The experiments were conducted on M41V/L/T cells 18 days after switching. **D:** mRNA expression levels of *IFI27, MX1,* and *CXCL10* decreased after treatment with GSK’872. The experiments were conducted on M41V/L/T cells 18 days after switching. **E:** mRNA expression levels of *IL1B* and *RNASE1* decreased after treatment with GSK’872. The experiments were conducted on M41V/L/T cells 22 days after switching. *\*P* < 0.05*, **P* < 0.01*, ***P* < 0.001, ns: not significant. Data are shown as means ± SD. *P* values were determined using a two-sided Kruskal-Wallis test, followed by Dunn’s post hoc test for multiple group comparisons (**A, B**), and a two-sided t-test (**C-E**). Data were obtained from two independent clones per genotype and pooled from two to four independent repeat experiments (**A-E**).

Further highlighting the central role of RIPK3, GSK’872 suppressed Tet-Off–induced *XBP1s* upregulation (Figure 7C). Additionally, GSK’872 almost completely abolished *IL1B* induction and *RNASE1* expression, as well as ISGs (*IFI27*, *MX1*, *CXCL10*), across all genotypes (Figure 7D-E). Thus, the RIPK3 inhibitor GSK’872 suppressed all pathological features of VS cell lines, including cell death, UPR, inflammation, and *RNASE1* expression.

## Discussion

In the current study, DP of patients identified RNASE1 as a potential biomarker of VS. Upon establishing human monocytic cell lines harboring *UBA1* mutations, we identified RIPK3 as a key molecule involved in various aspects of abnormalities in VS cells. Our data also demonstrated that the type of *UBA1* mutation defines the genotype-dependent phenotypic characteristics.

We propose that RNASE1 could function as a biomarker reflecting disease activity in VS. Although the disease specificity of RNASE1 remains unclear, its strong correlation with VEXASCAF suggests potential utility in assessing disease activity in patients with VS. C-reactive protein (CRP) is included in the VS remission criteria by the FRENVEX group;^16^ however, its levels can be affected by IL-6 inhibitors. Reports on the efficacy of IL-6 inhibition in VS are inconsistent.^41^ In some of our patients, despite normalization of CRP levels under IL-6 inhibitor treatment, RNASE1 remained elevated and inflammatory symptoms worsened, necessitating increases in PSL dosage. Therefore, RNASE1 may reflect disease activity more accurately and has the potential to serve as a superior biomarker to CRP.

RNASE1 is known to be highly expressed in the pancreas and vascular endothelial cells.^42^ We found that *RNASE1* is also prominent in immune cells, particularly monocytes, of patients with VS. Our data on the effects of the RIPK3 inhibitor in human monocytic VS cell lines indicate that RIPK3 activation ultimately induces *RNASE1* expression. Reduced ubiquitination due to *UBA1* mutations likely triggers the RIPK1-RIPK3 pathway, as RIPK1 kinase activity is negatively regulated by ubiquitination.^43–45^ This may be related to the elevated plasma levels of RIPK1 and RIPK3 in patients with VS.^31^ Consistent with our notion, a recent report has shown that VS mutations impair RIPK1 ubiquitination, thereby triggering inflammatory cell death.^39^

*RNASE1* gene expression is induced through JNK signaling in endothelial cells.^46^ We initially hypothesized that DAMPs released during necroptosis might contribute to inducing *RNASE1* via JNK in VS cells; however, this hypothesis was ultimately unfounded. We speculate that JNK may be activated by inflammatory cytokines and UPR,^47^ both of which are dependent on RIPK3, leading to the induction of RNASE1. Our results demonstrated the trace of JNK activation, namely, chromatin opening at the AP-1 elements, in VS cells. Supporting our notion, inflammation and UPR preceded the induction of *RNASE1* expression. Regarding the function of RNASE1, our knockdown experiments in VS cell lines did not detect notable effects; further investigation is warranted to determine whether RNASE1 is implicated in VS pathogenesis in vivo.

Multiple studies, including the current study, have demonstrated a strong UPR in VS cells.^1,25,31,48^ This has been attributed to the accumulation of misfolded proteins resulting from insufficient ubiquitination. However, our finding that RIPK3 activation influenced UPR suggests the possibility that increased sensitivity to endoplasmic reticulum (ER) stress also contributes to augmenting UPR. Reportedly, MLKL, activated by RIPK1 and RIPK3, translocates to the ER and damages the ER membrane during necroptosis.^49^ Another function of RIPK3, independent of necroptosis and UPR, is to promote IFN production by enhancing STING pathway signaling.^50^ In our study, we observed that RIPK3 activity is involved in the induction of ISGs.

Regarding the cell death pathways in VS, our results demonstrated that the activation of RIPK3 triggers necroptosis in VS cells. However, Annexin V^+^ cells likely contained not only early necroptosis but also apoptosis, possibly via ER stress.^51^ Monocytes from patients with VS exhibit elevated expression of genes related to PANoptosis, which includes pyroptosis.^31^ Notably, despite the induction of *IL1B*, we did not detect IL-1β secretion from our VS cell lines unless the NLRP3 activator nigericin was added. Thus, inflammasome activation and pyroptosis do not appear to be intrinsically executed within the monocyte population and may require additional extrinsic factors.

We propose that RIPK3 is an attractive target for treating patients with VS. RIPK1/3 inhibitors have been shown to reduce inflammation and mortality in systemic inflammatory response syndrome.^52^ Clinical trials of RIPK1 inhibitors for several inflammatory and neurodegenerative diseases were discontinued owing to insufficient efficacy.^53^ A phase 2 trial evaluating a RIPK1 inhibitor for ulcerative colitis is ongoing (NCT05588843). As RIPK3 is known to mediate necroptosis via both RIPK1-dependent and independent pathways,^33^ RIPK3 inhibitors may offer broader therapeutic potential. However, the development of RIPK3 inhibitors in mouse models has been unsuccessful. High-dose treatment of mouse cells with inhibitors for RIPK3 kinase activity *in vitro* induced apoptosis, and mice carrying a kinase-dead RIPK3 mutant (D161N) also underwent RIPK1-dependent spontaneous apoptosis.^54^ In both cases, the apoptosis was due to the dimerization of RIPK3 through its kinase domain.^55^ Therefore, RIPK3 inhibitors that abolish kinase activity without inducing dimerization warrant further consideration.

We demonstrated that *UBA1* mutations drive genotype-dependent phenotypes. In our VS cell lines, inflammation and cell death were most pronounced in M41V cells and least pronounced in M41L cells, mirroring clinical associations; M41V predicts poorer outcomes, whereas M41L correlates with better prognosis.^11–13^ The observed differences in reduced ubiquitination among genotypes suggest that genotype-specific effects contribute to disease pathogenesis at the upstream level.

Our results showing high VEXASCAF scores in the M41L group compared with those in the M41V and M41T groups have implications (Figure 3E). Although M41L cells displayed low pathogenicity, we anticipate that a substantial burden of mutant cells (high VAFs) in patients with M41L may lead to clinical symptoms (high VEXASCAF scores). In contrast, highly pathogenic M41V and M41T cells cause symptoms even with low VAFs. Similarly, the highest RNASE1 expression was observed in patients with M41L. However, cell lines harboring M41L did not exhibit *RNASE1* induction under nutrition-rich conditions. This discrepancy might be attributed to high VAFs in patients with M41L, who are often diagnosed only after the onset of severe symptoms in Japan.

These findings indicate the possible utility of patient stratification based on *UBA1* genotype, VAF, and RNASE1 expression. For instance, individuals with M41V or M41T may require intensive therapeutic intervention, even if the VAF and RNASE1 levels are low. In contrast, patients harboring M41L may require less intervention until the VAF and RNASE1 levels are elevated.

In conclusion, our study identified RNASE1 and RIPK3 as a potential biomarker of disease activity and an attractive therapeutic target in VS, respectively, and provides a basis for genotype-guided treatment strategies. Further investigations of the RIPK3-RNASE1 axis *in vivo* using mouse models, as well as clinical samples from real-world registry studies with larger sample sizes and extended follow-up periods, will be a key direction.

## Methods

### Biological samples from humans

Patients with suspected vacuole, E1 enzyme, X-linked, autoinflammatory, somatic (VEXAS) syndrome (VS) and patients with Behçet’s disease at Yokohama City University Hospital were enrolled in the study. In addition, healthy volunteers were recruited as controls. All patients with suspected VS had a score of ≥3, which predicted *UBA1* mutation positivity, as proposed by our group.^21^ All enrolled patients with suspected VS underwent *UBA1* genotyping using genomic DNA extracted from whole blood samples at the time of study enrollment. Peripheral blood and plasma were collected and stored in the Yokohama City University (YCU) biobank at −80°C until analysis. Total RNA was purified from peripheral blood stabilized in PAXgene Blood RNA Tubes (Cat. No. 762165; BD Biosciences, San Jose, CA, USA) using a PAXgene Blood miRNA Kit (Cat. No. 763134; QIAGEN, Venlo, The Netherlands), according to the manufacturer’s protocol. Peripheral blood mononuclear cells (PBMCs), monocytes, and neutrophils were isolated using the EasySep Direct Human PBMC/Monocyte/Neutrophil Isolation Kit (Cat. Nos. 19654/19669/19666; STEMCELL Technologies, Vancouver, BC, Canada). Human peripheral PBMCs, monocytes, and neutrophils were stained with antibodies (Supplemental Table 7) to confirm their purity after cell fractionation. Flow cytometry was performed using FACSCelesta (BD Biosciences), and the data were analyzed using FlowJo software (BD Biosciences). The purity of each immune cell type was approximately 90%.

### UBA1 genotyping

For *UBA1* genotyping, targeted amplicon deep sequencing (TAS) with a depth of over 3,000×, covering all exons of the *UBA1* gene, was performed at the Kazusa DNA Research Institute (Kisarazu, Chiba, Japan). DNA samples were prepared using the NEXTFLEX Rapid XP V2 DNA-seq Kit (Cat. No. NOVA-5149-23; Revvity, Waltham, MA, USA) and sequenced on the NextSeq platform (Illumina, San Diego, CA, USA) with 150-nt paired-end reads. Sequencing data were processed using fastp, BWA, Mutect2 (GATK v4.4.0.0), VarDict v1.0, and LoFreq v2.1.5, with the human reference genome GRCh38 (hg38). Mutations were retained if they met the following criteria: (1) allele frequency <1% in the Japanese population; (2) variant allele frequency >2%; and (3) passed the Mutect2 variant calling filter. Complementary DNA (cDNA) was synthesized from peripheral blood stabilized in PAXgene Blood RNA Tubes using the PrimeScript II 1st Strand cDNA Synthesis Kit (Cat. No. 6210A; Takara Bio, Tokyo, Japan) according to the manufacturer’s instructions. The synthesized cDNA was then used for TAS to quantify the VAF at each time point (Supplemental Table 8).

### Cell culture

Human monocytic cell line THP-1 (NIHS [JCRB], JCRB0112.1) was cultured in RPMI (Cat. No. 30263-95; Nacalai Tesque, Kyoto, Japan) with 10% fetal bovine serum (FBS), tetracycline-free, sterile-filtered (Cat. No. S181T-500; Biowest, Nuaillé, France), 100 U/mL penicillin, and 100 μg/mL streptomycin (PS) (Cat. No. 4987222637756, 4987222002929; Meiji Seika Pharma, Tokyo, Japan). HEK293T cells (RIKEN BRC CELL BANK, RCB2202) for lentivirus and retrovirus production were cultured in Dulbecco’s modified Eagle’s medium (DMEM; Cat. No. 08458-16; Nacalai Tesque, Kyoto, Japan) with 10% FBS (Cat. No. S1400-500; Biowest) and PS. NIH3T3 cells (CRL-1658; ATCC) for viral titration were cultured in DMEM supplemented with 10% Donor Bovine Serum (Cat. No. S0600; Biowest) and 1% PS. All cell lines were maintained in humidified incubators at 37°C and 5% CO_2_.

### Reverse transcription-quantitative PCR (RT-qPCR)

Total RNA was extracted from VS cell lines using an RNeasy Micro Kit (Cat. No.74004; QIAGEN) or a NucleoSpin RNA kit (Cat. No. U0955C; Takara Bio). RNA concentration was assessed using DS-11 (DeNovix, Wilmington, USA). cDNA was synthesized from total RNA using a PrimeScript RT Reagent Kit (Cat. No. RR037B; Takara Bio) according to the manufacturer’s instructions. The cDNA was mixed with the Thunderbird SYBR qPCR Mix (Cat. No. QPS-201; Toyobo, Osaka, Japan) and subjected to RT-qPCR on a CFX96 Real-Time System (Bio-Rad) using CFX Manager software. As per the manufacturer’s instructions, a typical two-step PCR protocol was performed (1 min at 95°C followed by 40 cycles of 15 s at 95°C and 45 s at 60°C). Gene-specific primers used in this study, designed using Primer-BLAST (NCBI), are listed in Supplemental Table 11. The data were normalized to the expression levels of glyceraldehyde 3-phosphate dehydrogenase (*GAPDH*).

### Plasmid construction

The cDNA of human UBA1 isoform b (RefSeq accession no. NP_003325.2) was obtained by reverse transcription polymerase chain reaction (RT-PCR) using RNA from THP-1 cells. The pRetroX-TRE3G-UBA1b construct was generated by inserting *UBA1b* cDNA into the pRetroX-TRE3G vector (Cat. No. 631188; Takara Bio).

### Lentiviral and retroviral vector preparation

HEK293T cells were seeded at 8×10^5^ cells/well into 6-well plates with 2.7 mL D-MEM 24 h before transfection. Lentiviral vector for Cas9 transduction was produced by transfecting HEK293T cells with 4.6 μg of pKLV2-EF1a-Cas9Bsd-W (lentiviral expression plasmid), 1.25 μg of pMDLg/pRRE (packaging plasmid), 0.6 μg of pRSV-Rev (packaging plasmid), and 0.75 μg of pMD2.G (envelope plasmid) (Cat. Nos. 68343, 12251, 12253, and 12259, respectively; Addgene, Cambridge, MA, USA) using 5 μL of PEI (2 μg/μL) (Cat. No. 24765; Polysciences, Warrington, PA, USA) in 300 μL Opti-MEM I reduced serum medium (Cat. No. 31985062; Thermo Fisher Scientific, Waltham, MA, USA). Retroviral vectors for Tet-On inducible *UBA1b* expression were produced by transfecting HEK293T cells with 2.0 μg of either pRetroX-TRE3G-UBA1b or pRetroX-Tet3G plasmid encoding the reverse tetracycline transactivator (Cat. No. 631188; Takara Bio), 1.5 μg of the packaging plasmid pMD.ogp (a gift from H. Xiong, Jining Medical University, Jining, Shandong, China), and 0.5 μg of the envelope plasmid pVSV-G (Clontech, Palo Alto, CA, USA) using 5 μL of PEI in 300 μL Opti-MEM I. Retroviral and lentiviral supernatants were collected 24 h post-transfection, filtered through a 0.45 μm syringe filter, and either used immediately for transduction or stored at −80°C. Viral titers were measured in NIH3T3 cells because HEK293T cells are resistant to neomycin (G418).

### Lentiviral and retroviral transduction

For lentiviral Cas9 transduction, 2 × 10^5^ THP-1 cells in 100 μL of culture medium were transduced in 24-well plates with 300 μL of lentiviral supernatant and 4 μg/mL polybrene (Cat. No. H9268-5G; Sigma-Aldrich, St Louis, MO, USA). After a 5-h incubation, 1 mL of fresh medium (without polybrene) was added, and the cells were cultured for an additional 24 h before being treated with 10 µg/mL blasticidin (Cat. No. ant-bl-05; InvivoGen, San Diego, CA, USA) to select transduced cells.

For retroviral transduction for Tet-On inducible *UBA1b* expression, 1 × 10^5^ Cas9-transduced THP-1 cells in 100 μL of culture medium containing doxycycline (Dox, final concentration: 1 µg/mL) were transduced in 24-well plates with 300 μL of each retroviral supernatant (multiplicity of infection [MOI] = 0.2) and 4 μg/mL polybrene. The plates were centrifuged at 1,200*×g* for 90 min at 32°C (spinoculation). After 5 h, the cells were washed and resuspended in a fresh medium lacking polybrene, followed by incubation for 24 h. The selection was performed using 1 mg/mL G418 (Cat. No. 04727878001; Roche, Basel, Switzerland) and 0.5 µg/mL puromycin (Cat. No. ant-pr-1; InvivoGen).

### Generation of VS model cell lines using CRISPR-Cas9-based homology-directed repair (HDR) mutation knock-in

The Cas9-transduced THP-1 cells harboring Tet-On inducible *UBA1b*, described above, were cultured with 1 µg/mL Dox to sustain exogenous *UBA1b* expression. Single-cell clones were isolated by limiting dilution, and the clone that demonstrated the most efficient depletion of exogenous UBA1b under Tet-Off conditions, determined by RT-qPCR, was selected for further experiments. The mutations in *UBA1* were introduced using the CRISPR-Cas9-mediated homology-directed repair (HDR) technique. HDR modulator RS-1 (7.5 µM) (Cat. No. R9782; Sigma-Aldrich) and NHEJ modulator SCR7 pyrazine (40 µM) (Cat. No. SML1546; Sigma-Aldrich) were added to the medium 4 h before electroporation. The cells were electroporated with single guide RNA (sgRNA) and donor oligo-DNA (Supplemental Table 9) using Nucleofector 2b (AAB-1001, Lonza, Cologne, Germany) and Cell Line Nucleofector Kit V (Cat. No. VCA-1003; Lonza) to knock-in the *UBA1* mutations (M41V, M41T, M41L) and a silent single-base mutation near c.121A (“unmutated” control) into *UBA1*. Twenty-four hours after electroporation, genomic DNA was isolated and assessed for *UBA1* mutations. Single-cell clones were further isolated by limiting dilution and screened for *UBA1* mutant clones. Sanger sequencing was performed to confirm that the mutations were incorporated as designed. Consequently, multiple clones were obtained for each mutation and unmutated control. Dox was removed from the medium during downstream assays. For the phenotypic analysis, two or more clones with the closest doubling times were selected for each genotype. The established VS cell lines were maintained with 1 μg/mL Dox and cryopreserved using CELLBANKER 1 (Cat. No. CB011; ZENOAQ, Fukushima, Japan).

### Enzyme-linked immunosorbent assay (ELISA)

Human serum and plasma ribonuclease 1 (RNASE1) levels were measured using ELISA kits purchased from Sino Biological Inc. (Cat. No. SEK13468; Beijing, China). THP-1 cells were seeded at a density of 1 × 10⁶ cells/mL in 96-well culture plates, incubated for 24 h, and then stimulated with 10 µM nigericin (Cat. No. 149-07261; WAKO, Osaka, Japan) for 2 h. Culture supernatants were collected and analyzed using commercially available ELISA kits; IL-18 levels were measured using a kit from R&D Systems (Cat. No. DY318-05; Abingdon, UK), and IL-1β levels were measured using the kit from BioLegend (Cat. No. 437004; San Diego, CA, USA).

### Annexin V/propidium iodide assay

Cell death was assessed using Annexin V/propidium iodide staining (Cat. No. 640914; BioLegend), according to the manufacturer’s protocol. Stained cells were analyzed by flow cytometry using a FACSCelesta. The Annexin V-positive rate (%) was calculated as follows: (sum of Annexin V-positive cells) / (total cell number) × 100.

### Immunoblotting analysis

Peripheral blood monocytes were isolated using the EasySep Direct Human Monocyte Isolation Kit. To prevent degradation or modification of the proteins, protease inhibitor phenylmethylsulfonyl fluoride (Cat. No. 27327-52, Nacalai Tesque), phosphatase inhibitors cypermethrin (Cat. No. ab141018, Abcam, Cambridge, MA, USA), and okadaic acid (Cat. No. sc-3513A, Santa Cruz Biotechnology, Santa Cruz, CA, USA) were added into peripheral blood shortly after blood sampling. For nuclear and cytoplasmic fractionation, cells were processed using Buffer A (HEPES, pH 7.4; KCl; MgCl₂; protease inhibitor cocktail; phosphatase inhibitor cocktail; and NP-40). Cells were resuspended in Buffer A containing 0.5% NP-40, followed by incubation on ice for 5 min. Samples were centrifuged at 1,000 × g for 5 min at 4°C, and the supernatant was collected as the cytoplasmic fraction. Nuclear pellets were washed once with detergent-free Buffer A and centrifuged under the same conditions. After removal of the wash buffer, nuclear proteins were extracted by resuspending the pellets in RIPA buffer with gentle pipetting. Whole-cell lysates were prepared by lysing cells in RIPA buffer on ice for 30 min in the presence of protease and phosphatase inhibitors (Cat. Nos. 11836170001 and 04906837001; Roche). Lysates were collected by centrifugation at 17,700 × g for 15 min. The lysates were analyzed by performing sodium dodecyl sulfate-polyacrylamide gel electrophoresis. After proteins were transferred onto PVDF membranes (Cat. No. 1704272; Bio-Rad, Richmond, CA, USA), the membranes were blocked with 5% non-fat dry milk for 30 min at room temperature, followed by incubation with primary antibodies (Supplemental Table 10) overnight at 4°C. After washing, the membranes were incubated with the appropriate secondary antibodies (Supplemental Table 10). The protein bands were detected using the standard chemiluminescence method with ECL Prime (Cat. No. RPN2232; Cytiva, Uppsala, Sweden) and imaged using a Fusion Solo S system (Vilber Lourmat, Paris, France).

### Wright–Giemsa stain

In brief, 2 × 10^4^ cells were harvested from culture and spun down onto glass slides (Cat. No. S2215; Matsunami Glass, Osaka, Japan) in a CytoSpin 4 Cytocentrifuge (A78300003; Thermo Fisher Scientific) at 500 rpm for 1 min. Cells were immediately stained with Wright–Giemsa (Cat. No. 15022; Muto Pure Chemicals, Tokyo, Japan) according to the manufacturer’s protocol. Images were acquired using a MICA microscope (Leica Microsystems, Wetzlar, Germany) equipped with a ×63 objective lens.

### Measurement of XBP1 splicing

Total and spliced *XBP1* mRNA levels were quantified using RT-qPCR with specific primers (Supplemental Table 11). The percentage (%) of spliced XBP1 was calculated as (spliced XBP1 / total XBP1) × 100.

### shRNA-mediated gene silencing of *RNASE1*

Lentiviral vectors containing one of three *RNASE1*-specific shRNA constructs or a scrambled shRNA as a negative control (Supplemental Table 12) were purchased from Vector Builder Japan (Kanagawa, Japan). Lentiviral transduction was performed on both VS and unmutated cell lines at an MOI of 2 with 4 µg/mL polybrene. After 5 h, fresh medium without polybrene was added, and the cells were cultured for an additional 24 h before selection with 400 µg/mL hygromycin (Cat. No. ant-hg-1; InvivoGen) to enrich the transduced cells. Drug selection was continued for 7 days. *RNASE1* knockdown efficiency was quantified using RT-qPCR. The established knockdown cells were maintained using 20 ng/mL Dox and cryopreserved in CELLBANKER 1.

### Inhibitor assay

To evaluate the sensitivity of VS cell lines to the UBA1 inhibitor, cells were treated for 24 h with either DMSO or TAK-243 (Cat. No. S8341; Selleck Chemicals, Houston, TX, USA) at concentrations of 10 nM and 1,000 nM. To assess the involvement of cell death pathways in VS cell lines, cells were cultured under Tet-On/Off conditions in medium supplemented with 10 µM each of the pan-caspase inhibitor Z-VAD-FMK (Cat. No. 3188-v; Peptide Institute, Inc., Osaka, Japan) and 10 µM RIPK3 inhibitor GSK’872 (Cat. No. S8465; Selleck Chemicals). To clarify the contribution of IRF, AP-1, and necroptosis to the pathological state of VS, M41V cell lines were cultured under Tet-On/Off conditions in medium supplemented with 30 µM RIPK1 inhibitor Necrostatin-2 racemate (Nec-1s; Cat. No. S8641; Selleck Chemicals), 10 µM RIPK3 inhibitor GSK’872, 10 µM JNK inhibitor SP600125 (Cat. No. S1460; Selleck Chemicals), 10 µg/mL anti-IFNAR1 antibody anifrolumab (Cat. No. A2460; Selleck Chemicals), or 10 µg/mL human IgG1 isotype control (Cat. No. A2051; Selleck Chemicals).

### Caspase activity assay

To confirm the effect of Z-VAD-FMK, VS cell lines (5 × 10^4^ cells/well) were seeded in 96-well plates, followed by the addition of Caspase-Glo 1 (Cat. No. G9951; Promega, Madison, WA, USA). The luminescence intensity (LI) was measured, and the Relative Luminescence Units were calculated by normalizing the LI values to the background signal.

### Transwell assay

To determine whether damage-associated molecular patterns released from VS cell lines induce *RNASE1* expression in a cell-extrinsic manner, VS cell lines cultured for 14 days under Tet-Off conditions were seeded onto the upper side of a transwell insert with a 3.0-μm pore membrane (Cat. No. 3414; Corning, NY, USA), and unmutated cells were placed in the lower chamber for coculture over 9 days. In a similar setup, M41V cells cultured under Tet-Off conditions for 24 days were seeded onto the upper side of a transwell insert, while M41V cells cultured under Tet-Off conditions for 14 days were seeded in the lower chamber, and the cells were then cocultured for 8 days.

### Bulk RNA sequencing analysis

For PAXgene-stabilized whole blood, total RNA was extracted using a PAXgene Blood miRNA Kit (Cat. No. 763134; QIAGEN) according to the manufacturer’s protocol. Total RNA from whole blood and a part of unmutated cell line samples (Supplemental Figure 8B) were subjected to RNA quality checks, globin mRNA depletion (for whole blood RNA), library synthesis, and sequencing by Takara Bio, Inc.

For PBMCs, monocytes, and neutrophils, as well as for VS and unmutated cell lines, total RNA was extracted using the RNeasy Micro Kit (Cat. No. 74004; QIAGEN). RNA was assessed using a 4150 TapeStation system (Agilent Technologies, Foster City, CA, USA) with a High-Sensitivity RNA ScreenTape, Sample Buffer, and Ladder (Cat. Nos. 5067-5579, 5067-5580, and 5067-5581, respectively; Agilent Technologies). All RNA samples had an RNA integrity number >8.9. Libraries were prepared from 50 to 250 ng of total RNA using the SureSelect XT HS2 mRNA Library Preparation Kit (Agilent Technologies) according to the manufacturer’s instructions. The final DNA library was assessed using the 4150 TapeStation system with High Sensitivity D1000 ScreenTape and Reagents (Cat. Nos. 5067-5584 and 5067-5585, respectively; Agilent Technologies). The concentrations of individual libraries were measured by qPCR using a KAPA Library Quantification Kit (Cat. No. KK4824; Nippon Genetics, Tokyo, Japan). Pooled libraries were sequenced using the NextSeq 500 System (Illumina) with the NextSeq 500/550 High Output Kit v2 (75 cycles) (Cat. No. 20024906; Illumina). Raw sequence data in the FASTQ format were obtained using bcl2fastq (ver. 2.20.0.422). Public RNA-seq read data were retrieved using the parallel-fastq-dump command (ver. 0.6.7).

The FASTQ files containing paired-end reads were assessed using FastQC (ver. 0.11.9) or preprocessed to trim adapter sequences and filter out low-quality reads using Fastp (ver. 0.23.4). The reads were then aligned to the GRCh38 (hg38) human reference genome using the STAR aligner (ver. 2.7.10a or 2.7.11b). Aligned reads were processed using StringTie (ver. 2.2.1) with Gencode annotations (v39 or v45) to obtain raw and normalized count data. R software (ver. 4.3.2) was used to analyze and visualize the data. Differential expression analyses were performed using the DESeq2 package (ver. 1.42.0). A principal component analysis was performed using the prcomp function. For analyzing genes whose expression correlated with VEXASCAF, normalized counts obtained by the cpm function were z-score scaled by the genescale function in the genefilter package (ver. 1.84.0) and then subjected to Spearman’s rank correlation coefficient calculations using the rcorr function from the Hmisc package (ver. 5.1.1). *P*-values adjusted using the Benjamini-Hochberg method were calculated using the p.adjust function. Volcano plots were generated using the EnhancedVolcano package (ver. 1.20.0). GSEA of the MSigDB hallmark or necroptosis response^34^ gene sets was performed using the GSEA function of the clusterProfiler package (ver. 4.10.0).

### Single-cell RNA sequencing (scRNA-seq) analysis

Isolated PBMCs and neutrophils were combined in a ratio of 4:1 and subjected to the scRNA-seq library preparation using the Chromium Controller (10x Genomics, Pleasanton, CA, USA) with Chromium Next GEM Single Cell 3ʹ GEM, Library & Gel Bead Kit v3.1, Chip G Single Cell Kit, and Single Index Kit T Set A (Cat. nos. 1000121, 1000127, and 1000213, respectively; 10x Genomics) according to the manufacturer’s protocol. The final DNA library was assessed using the 4150 TapeStation system with High Sensitivity D1000 ScreenTape and Reagents. The concentrations of individual libraries were measured using qPCR with the KAPA Library Quantification Kit. Sequencing was performed in the paired-end mode (read1:28 bp; read2:91 bp) using a NextSeq 500 sequencer (Illumina) with a NextSeq 500/550 High Output Kit v2.5 (150 cycles) (Cat. No. 20024907; Illumina). Public scRNA-seq read data were retrieved using the parallel-fastq-dump command (ver. 0.6.7).

The resulting scRNA-seq reads were processed using the CellRanger pipeline (ver. 7.1.0) against the refdata-gex-GRCh38-2020-A reference genome in 3’ v3 chemistry mode (10x Genomics). Subsequent analyses were performed using the Seurat R package (ver. 5.0.2). Cells were filtered based on the thresholds of a minimum unique molecular identifier count of 100 (for the detection of neutrophils), a maximum number of detected genes of 6,000, and a maximum percentage of mitochondrial genes of 10%. Data were normalized using the NormalizeData function and integrated using the IntegrateData function for reference-based integration. Canonical correlation analysis was used as a reduction method to identify the anchors using the FindIntegrationAnchor function. Cluster identification of cells was performed by a shared nearest neighbor modularity optimization-based algorithm using the FindNeighbors and FindClusters functions. The UMAP and violin plots were generated using the DimPlot and VlnPlot functions, respectively. Gene expression and density in the UMAP plots were visualized using the FeaturePlot_scCustom and Plot_Density_Custom functions, respectively, in the scCustomize package (ver. 3.0.1). Statistical comparisons between the cell groups were performed using the Wilcoxon rank-sum test.

### ATAC-seq analysis

Preparation of the ATAC-seq library from VS or unmutated cell lines was performed as described previously.^56,57^ Briefly, 50,000 cells were incubated with ATAC-seq resuspension buffer (RSB) containing 0.1% NP-40, 0.1% Tween-20, and 0.01% digitonin for 3 min on ice to isolate the nuclei. Subsequently, nuclei were incubated with a transposition mix (1× TD buffer, 100 nM transposase [TDE1; Cat. no. 20034198; Illumina], 33% 1× PBS [33% of the total volume of the transposition mix was 1× PBS], 0.01% digitonin, 0.1% Tween-20, and 5% water) for 30 min at 37°C with shaking at 900 rpm. The tagged DNA was purified using the MinElute Reaction Cleanup Kit (Cat. No. 28204; QIAGEN). Purified tagmented DNA was amplified using a NEBNext High Fidelity 2× PCR Master Mix (Cat. No. M0541L; New England Biolabs, Ipswich, MA, USA) with indexed primers. Libraries were purified using the Agencourt AMPure XP Kit (Cat. No. A63881; Beckman Coulter, Brea, CA, USA). The final DNA library was assessed using the 4150 TapeStation system with High Sensitivity D1000 ScreenTape and Reagents. The pooled libraries were sequenced using a NextSeq 500 System with a NextSeq 500/550 High Output Kit v2 (75 cycles). Raw sequence data in the FASTQ format were obtained using bcl2fastq (ver. 2.20.0.422).

The FASTQ files containing paired-end reads were assessed using FastQC (ver. 0.12.1). The reads were then aligned to the GRCh38 (hg38) human reference genome using the bowtie2 aligner (ver. 2.5.1). Aligned reads were processed using samtools (ver. 1.17) and picard (ver. 3.0.0) to obtain reads de-duplicated and filtered for high quality (MAPQ ≥ 30). Peak calling was performed using MACS3 (version 3.0.3) with the following parameters: --nomodel, --shift −37, --extsize 74, and --keep-dup all. Consensus peaks among replicates were extracted using bedtools intersect (version 2.30.0). The consensus peaks for all sample groups were merged using bedops (version 2.4.41) to generate an ATAC-seq peak catalog. The read counts for each peak in the catalog were measured using featureCounts (ver. 2.0.6). Scale factors were calculated as 1 / (NormFactor × LibSize / 1,000,000), where NormFactor was obtained using the calcNormFactors function in edgeR (ver. 4.0.12) and LibSize is the raw library size. The bamCoverage function in deepTools (ver. 3.5.6) was executed with -of parameter set to bigwig and --scaleFactor parameter set to the scale factor values. The resulting bigwig and peak catalog files were used to visualize the ATAC-seq data in the UCSC Genome Browser (https://genome.ucsc.edu/cgi-bin/hgGateway). R software (ver. 4.3.2) was used for the subsequent analyses. Peaks were annotated using the annotatePeak function in the ChIPseeker package (ver. 1.38.0). The GC content of the peak regions was calculated using the letterFrequency function of the Biostrings package (ver. 2.70.2) for GC-aware normalization. A differential accessibility analysis was performed using edgeR (ver. 4.0.12) and EDASeq (ver. 2.36.0) packages. The GSEA plots were generated using the EnhancedVolcano package (ver. 1.20.0). Motif analysis was performed using Homer software (ver. 5.1), with the marge package (ver. 0.0.4.9999) to facilitate the query construction. *De novo* motifs were obtained using the findMotifsGenome.pl function with “-size given” parameter.

### Ethical approval

This study was approved by the Yokohama City University Certified Institutional Review Board (Approval ID: F22070005). Informed consent was obtained from all participants or their legal guardians. The protocol for the prospective VS registry has been registered in the UMIN database (UMIN000053426).

### Statistical analysis

Data analyses were performed using R software (https://www.r-project.org/). Non-normally distributed data were compared using the Mann–Whitney U test. Normally distributed data were compared either using Student’s *t*-test or Welch’s *t*-test. The Kruskal–Wallis test was used to compare two or more independent samples. Spearman’s rank-order correlation was used for the correlation analysis. Statistical significance was set at *P* < 0.05.

## Data availability

The bulk RNA-seq, scRNA-seq, and ATAC-seq data obtained in this study are available in the Gene Expression Omnibus (GEO) database (https://www.ncbi.nlm.nih.gov/geo/) under the following accession numbers: GSE298044 and GSE313020 for bulk RNA-seq of cell lines, GSE298045 and GSE313023 for bulk RNA-seq of peripheral blood, GSE298238 for bulk RNA-seq of monocytes, GSE298354 for scRNA-seq of PBMCs, and GSE298043 for ATAC-seq of cell lines. In GSE298043 and GSE298044, M41V clones 3 and 4 correspond to M41V clones 1 and 2; M41T clones 2 and 3 to M41T clones 1 and 2; and M41L clones 2 and 3 to M41L clones 1 and 2, respectively, as referred to in this study. Public data were obtained from the GEO database: GSE160268 for bulk RNA-seq of monocytes from patients with VS and healthy controls (HC)^1^; GSE226642 for bulk RNA-seq of monocytes from CHIP carriers and non-carriers^26^; GSE249131 and GSE196052 for scRNA-seq of PBMCs^24^ and bone marrow hematopoietic cells^25^, respectively, from patients with VS and age- and sex-matched HC.

## Acknowledgments

We thank the patients and their families for providing their medical histories and specimens. We also thank Yuki Hirabayashi and Professor Satoshi Fujii of the YCU Biobank and YCU Hospital Clinical Laboratory for their cooperation in storing the patient specimens. We thank Noriko Tagata of Yokohama City University for her assistance with the experiments. Figures 1A, 3E, and 4A, Supplemental Figure 1, 14, and the visual abstract were prepared using BioRender (https://www.biorender.com/). Supplemental Figure 7 was prepared using Sanger sequencing traces from Chromas 2.6.6 DNA Sequencing Software (https://technelysium.com.au/wp/chromas/). Grammarly (https://www.grammarly.com/) and DeepL (https://www.deepl.com/) were used to improve the grammar of the manuscript. We thank Editage (https://www.editage.jp) for English language editing. This work was supported by the Japan Agency for Medical Research and Development (grant JP24ek0410107) and partly by a joint research fund from Nippon Shinyaku.

## Authorship

Contribution: K.H., T.B., and Y.K. designed the study; K.H., T.B., and G.R.S. performed the experiments; K.H., T.B., and G.R.S. analyzed the data; K.H., T.B., Y.K., and T.T. wrote the manuscript; S.A., Y.I., A.M., and O.O. provided key materials; H.N. and T.T. provided critical suggestions for the study; Y.K. supervised the deep phenotyping project; and T.B. supervised the disease model cell project.

