## Supplemental Data for "RIPK3-RNASE1 axis as a potential therapeutic and clinical monitoring target in VEXAS syndrome"

### Supplemental Tables

**Supplemental Table 1. Details of clinical and genetic features of patients with VS at the time of deep phenotyping entry.**

| ID | Age | Sex | Mutation | VAF (%) | O.P. | Symptoms at diagnosis | Treatment |
|---|---|---|---|---|---|---|---|
| 1 | 78 | M | Val | 11.3 | 388 | Fever, skin lesion, pulmonary lesion (pulmonary multinodular shadows), relapsing polychondritis, polyarthritis | PSL, TCZ |
| 2 | 71 | M | Leu | 87.5 | 123 | Fever, polyarthritis, skin lesion, MDS-MLD, relapsing polychondritis, giant cell arteritis | PSL, TCZ |
| 3 | 85 | M | Thr | 21.8 | 350 | Fever, skin lesion (Sweet's disease), MDS-MLD, pulmonary lesion (pulmonary infiltrate) | PSL, TCZ |
| 4 | 55 | M | Splice | 56.4 | 358 | Fever, myositis, skin lesion, MDS-MLD, pulmonary lesion (pulmonary multinodular shadows) | PSL |
| 5 | 71 | M | Leu | 45.6 | 367 | Fever, skin lesion (LCV) | - |
| 6 | 70 | M | Thr | 72 | 367 | Fever, skin lesion (LCV, Sweet's disease), conjunctival injection, ear chondritis | - |
| 7 | 71 | M | Other | 53.6 | 364 | Fever, skin lesion, MDS-SLD, polyarthritis, pulmonary lesion (GGO) | - |
| 8 | 88 | M | Val | 10.4 | 364 | Fever, polyarthritis, skin lesion (Sweet's disease), MDS-MLD, pulmonary lesion (organized pneumonia), RS3PE syndrome | PSL |
| 9 | 76 | M | Thr | 67.3 | 327 | Fever, polyarthritis, skin lesion, relapsing polychondritis, pulmonary lesion (GGO), DVT | PSL, TCZ |
| 10 | 72 | M | Leu | 86.1 | 277 | Fever, arthritis, skin lesions, relapsing polychondritis, scleritis | PSL |
| 11 | 77 | M | Leu | 51 | 101 | Fever, polyarthritis, skin lesion, relapsing polychondritis, thrombosis (cerebral, arterial), conjunctival injection | PSL, CyA |
| 12 | 80 | M | Leu | 64.2 | 147 | Polyarthritis, skin lesion, relapsing polychondritis, conjunctival injection | PSL |
| 13 | 73 | M | Thr | 62.8 | 31 | Fever, polyarthritis, relapsing polychondritis | - |

'Splice' refers to the c.118-1G>C mutation. 'Other' refers to the UBA1 p.Ala478Ser mutation.

VAF: variant allele frequency of *UBA1* mutations in whole blood data at entry.

O.P.: observation period, for which the term 'observation' indicates the number of days at the last observation point after entry.

MDS, myelodysplastic syndrome; LCV, leukocytoclastic vasculitis; GGO, ground-glass opacity; DVT, deep vein thrombosis; PSL, prednisolone; CyA, cyclosporin; TCZ, tocilizumab; VS, vacuole, E1 enzyme, X-linked, autoinflammatory, somatic (VEXAS) syndrome.

**Supplemental Table 2. Clinical manifestations of MDS in deep phenotyping participants.**

| Group | ID | MDS type | Age at MDS onset | Megakaryocyte | Myeloid | Erythroid | Cytoplasmic vacuolation in myeloid/erythroid progenitor cells | BM blast (%) | Cytogenetics | IPSS-R at MDS diagnosis | T.D. |
|---|---|---|---|---|---|---|---|---|---|---|---|
| VS | 2 | MDS-MLD | 69 | Micro-MgK, Multinucleation, Nuclear hypolobation | – | Karyorrhexis, Megaloblastoid changes | + | 0.2 | 46XY | 3 (low) | RBC |
| | 3 | MDS-MLD | 83 | Micro-MgK, Multinucleation | Decreased granules | – | + | 0.2 | 46XY | 2 (low) | – |
| | 4 | MDS-MLD | 54 | Micro-MgK, Multinucleation | Pseudo-Pelger-Huët, Decreased granules | – | + | 0.4 | 46XY | 2.5 (low) | RBC |
| | 7 | MDS-SLD | 68 | Micro-MgK | – | Megaloblastoid changes | – | 0.2 | 46XY, del(20q)(q11.2q 13.3) | 1 (very low) | – |
| | 8 | MDS-MLD | 83 | Micro-MgK, Multinucleation | Decreased granules | – | + | 1.6 | 46XY | 3 (low) | – |
| VLAD | 14 | MDS-MLD | 77 | Micro-MgK | Pseudo-Pelger-Huët, Decreased granules | Megaloblastoid changes | – | 0.2 | 47, XY, +8[17]/46XY[3] | 2 (low) | – |
| | 15 | MDS-MLD | 81 | Micro-MgK, Multinucleation | Hypogranulation, Pseudo-Pelger-Huët, Decreased granules | Megaloblastoid changes, Multinuclearity, Multinucleated erythroblast, Irregular nuclear contour | + | 3.6 | del(5)(q13q33), -7[4]/46XY[16] | 6 (high) | RBC |
| | 16 | MDS-MLD | 79 | Micro-MgK | Pseudo-Pelger-Huët, Decreased granules | Megaloblastoid changes | – | 1.0 | 47, XX, +8[16]/47,idem, del(11)(q?)[3]/4 6XX[1] | 2.5 (low) | PC |
| | 17 | MDS-U | 80 | Micro-MgK | – | – | – | 1.0 | 47, XY, +8[16]/48, idem, +mar[4] | 4 (intermediate) | RBC |
| | 18 | MDS-MLD | 81 | Segmented multinucleated megakaryocyte | – | Karyorrhexis | + | 1.6 | 47, XY, +8[19]/46XY[1] | 2 (low) | – |
| | 33 | MDS-U | 68 | Micro-MgK | – | – | – | 0.1 | 46XY | 1.5 (very low) | – |
| | 35 | MDS-SLD | 80 | Micro-MgK, Multinucleation | – | – | – | 0.1 | 45, X, -Y | 0.5 (very low) | – |
| | 40 | MDS-MLD | 66 | Non-lobulated nuclei | Pseudo-Pelger-Huët, Decreased granules | Karyorrhexis, irregular nuclear contour | – | 5.0 | 46XY | 2 (low) | – |

VS, vacuole, E1 enzyme, X-linked, autoinflammatory, somatic (VEXAS) syndrome; VLAD, VS-like autoinflammatory diseases; MDS-SLD, MDS with single lineage dysplasia; MDS-MLD, MDS with multilineage dysplasia; MDS-U: Unclassified MDS, MgK, megakaryocytes; IPSS-R, revised international prognostic scoring system; T.D., transfusion dependence; RCC, red cell concentrate; PC, platelet concentrate.

**Supplemental Table 3. Clinical characteristics of the two independent groups of patients with VS in deep phenotyping.**

| Patient samples | Discovery | Validation |
|---|---|---|
| Demographic characteristics of participants | | |
| Participants, n | 10 | 11 |
| Male sex, n (%) | 10 (100) | 11 (100) |
| Median age at study entry (range) | 71.5 (55–88) | 76 (55–88) |
| Genetic characteristics | | |
| p.Met41Val (c.121A > G), n (%) | 2 (20) | 2 (18.2) |
| p.Met41Thr (c.122T > C), n (%) | 3 (30) | 4 (36.4) |
| p.Met41Leu (c.121A > C), n (%) | 3 (30) | 3 (27.3) |
| Splice, n (%) | 1 (10) | 1 (9.1) |
| p.Ala478Ser, n (%) | 1 (10) | 1 (9.1) |
| Deep phenotyping | | |
| Samples, n | 38 | 41 |
| Clinical features | | |
| VEXASCAF -points (IQR) | 1 (0–2) | 1 (0–2) |
| Fever, n (%) | 1 (2.6) | 4 (9.8) |
| Ear chondritis, n (%) | 2 (5.3) | 1 (2.4) |
| Nose chondritis, n (%) | 1 (2.6) | 1 (2.4) |
| Lung symptoms, n (%) | 8 (21.1) | 1 (2.4) |
| Arthralgia, n (%) | 14 (36.8) | 8 (19.5) |
| Arthritis, n (%) | 8 (21.1) | 4 (9.8) |
| Vein thrombosis, n (%) | 1 (2.6) | 9 (22) |
| Dermatosis, n (%) | 8 (21.1) | 11 (26.8) |
| Eye inflammation, n (%) | 5 (13.2) | 1 (2.4) |
| Periorbital edema, n (%) | 0 (0) | 0 (0) |
| Vasculitis, n (%) | 4 (10.5) | 2 (4.9) |
| Laboratory data at entry | | |
| Hemoglobin, g/dL (IQR) | 8.8 (7.7–12.1) | 10.8 (7.4–12.5) |
| MCV, fL (IQR) | 109 (105–111) | 101 (97.8–108) |
| Leukocytes, $\times 10^9$/L (IQR) | 3.8 (2.6–5.0) | 3.1 (1.9–5.0) |
| Neutrophils, $\times 10^9$/L (IQR) | 2.8 (1.6–4.0) | 2.0 (1.0–3.5) |
| Monocytes, $\times 10^9$/L (IQR) | 0.20 (0.10–0.34) | 0.23 (0.13–0.32) |
| Platelets, $\times 10^3$/μL (IQR) | 82 (58–124) | 121 (78–156) |
| C-reactive protein, mg/L (IQR) | 1.0 (0.2–7.3) | 1.0 (0.2–4.0) |
| Glucocorticoids, n (%) | 33 (86.8) | 38 (92.7) |
| Median prednisone dose, mg/day (IQR) | 15 (10–40) | 17.5 (13–25) |
| Synthetic DMARDs, n (%) | 0 (0) | 2 (4.9) |
| Biologic DMARDs, n (%) | 26 (68.4) | 30 (73.2) |

'Splice' refers to the c.118-1G>C mutation. VS, vacuole, E1 enzyme, X-linked, autoinflammatory, somatic (VEXAS) syndrome; DMARDs, disease-modifying antirheumatic drugs; IQR, interquartile range; NA, not available. Laboratory data are presented as median values.

Synthetic DMARDs: tacrolimus, methotrexate, Biologic DMARDs: tocilizumab.

**Supplemental Table 4. Changes in clinical symptoms of patients with VS after intensified prednisolone treatment.**

| ID | Before increasing the PSL dosage | Four days after increasing the PSL dosage |
|---|---|---|
| X | Dyspnea on exertion, new lung infiltrates | Improved dyspnea, improved lung infiltrates |
| 8 | Fever, new lung infiltrates | Fever resolved, improved lung infiltrates |
| 4 | Fever, rash | Fever resolved, disappearance of rash |
| 2 | Difficulty in movement, worsening lung infiltrates, worsening rash | Mobility impaired, improved lung infiltrates, rash unchanged |

PSL, prednisolone; VS, vacuole, E1 enzyme, X-linked, autoinflammatory, somatic (VEXAS) syndrome.

**Supplemental Table 5. Clinical characteristics of patients with VS who underwent proteome and bulk RNA sequencing analysis for validation of RNASE1 expression**

| ID | Age | Sex | Mutation | Treatment | Symptoms at diagnosis |
|---|---|---|---|---|---|
| 4 | 55 | M | Splice | PSL | Fever, myositis, skin lesion, MDS-MLD, pulmonary lesion (pulmonary multinodular shadows) |
| 2* | 71 | M | Leu | PSL, TCZ | Fever, polyarthritis, skin lesion, MDS-MLD, relapsing polychondritis, giant cell arteritis |
| 8* | 87 | M | Val | PSL | Fever, polyarthritis, skin lesion (Sweet's disease), MDS-MLD, pulmonary lesion (organized pneumonia), RS3PE syndrome |
| 3 | 85 | M | Thr | PSL, TCZ | Fever, skin lesion (Sweet's disease), MDS-MLD, pulmonary lesion (pulmonary infiltrate) |
| X | 67 | M | Thr | PSL, TCZ, CY | Fever, polyarthritis, skin lesion (Sweet's disease), MDS-MLD, relapsing polychondritis, scleritis |

'*' refers to patients who underwent scRNA-seq. 'ID:X' is a patient who did not participate in deep phenotyping.

'Age' and treatment refer to information provided at the time of sample collection.

'Splice' refers to the c.118-1G>C mutation. PSL, prednisolone; TCZ, tocilizumab; CY, cyclophosphamide; VS, vacuole, E1 enzyme, X-linked, autoinflammatory, somatic (VEXAS) syndrome.

**Supplemental Table 6. Clinical data of patients with VS who underwent monocyte immunoblotting analysis**

| ID | Age | Sex | Mutation | VEXASCAF | Symptoms | Hemoglobin, g/dL | Platelets, $\times 10^3/\mu L$ | CRP, mg/L | T.D. | PSL, mg/day |
|---|---|---|---|---|---|---|---|---|---|---|
| 6 | 72 | M | Thr | 0 | − | 11.8 | 88 | 0.1 | − | 6.25 |
| 4 | 58 | M | Splice | 2 | Fever, subcutaneous abscess | 5.1 | 14 | 56.9 | + | 26 |

'Splice' refers to the c.118-1G>C mutation. CRP, C-reactive protein; T.D., transfusion dependence; PSL, prednisolone.

**Supplemental Table 7. Antibodies used in FACS analyses.**

| Antibody | Cat. No. | Source |
|---|---|---|
| Brilliant Violet 650 anti-human CD16 Antibody | 302041 | BioLegend |
| CD66b-FITC | IM0531U | Beckman Coulter |
| APC/Cyanine7 anti-human CD41 Antibody | 303716 | BioLegend |
| PE/Cyanine7 anti-human CD45 Antibody | 368532 | BioLegend |
| APC anti-human CD14 Antibody | 301808 | BioLegend |
| Brilliant Violet 510 anti-human CD19 Antibody | 302241 | BioLegend |
| Brilliant Violet 421 anti-human CD3 Antibody | 300434 | BioLegend |
| Brilliant Violet 605 anti-human HLA-DR Antibody | 307640 | BioLegend |
| PE anti-human CD235a (Glycophorin A) Antibody | 349106 | BioLegend |
| 7-Amino Actinomycin D | A9400-5MG | Sigma-Aldrich |

**Supplemental Table 8. Temporal dynamics of *UBA1* mutant VAF in patients with VS revealed by deep phenotyping analysis.**

| | *UBA1* mutant VAF from cDNA | | | | |
|---|---|---|---|---|---|
| ID | At entry | At 3 months | At 6 months | At 9 months | At 1 year |
| 1 | 28.2 | 42.0 | 39.5 | 16.7 | 18.8 |
| 2 | 87.8 | 88.6 | ND | ND | ND |
| 3 | 9.9 | 17.3 | 21.6 | 12.7 | 23.0 |
| 4 | 21.2 | 26.8 | 25.3 | 44.3 | 44.1 |
| 5 | 36.6 | 43.2 | ND | ND | ND |
| 6 | 55.8 | 46.4 | 30.3 | 19.5 | 15.9 |
| 7 | 27.6 | 35.6 | 22.3 | 37.6 | ND |
| 8 | 9.6 | 11.1 | 7.5 | 4.5 | 1.7 |
| 9 | 59.2 | 66.9 | ND | 83.0 | ND |
| 10 | 62.8 | ND | 82.6 | ND | ND |
| 11 | 52.5 | 42.9 | ND | ND | ND |
| 12 | 58.4 | ND | ND | ND | ND |
| 13 | 50.0 | ND | ND | ND | ND |

ND: Not determined.

**Supplemental Table 9. Oligonucleotides and sgRNA used in CRISPR/Cas9-based HDR.**

| | Name | Sequence |
|---|---|---|
| Oligo-DNA | VS (M41V) | 5′ -T*C*CACACCCCTCAGCCCGGCCTTGGCCCCACTTACAGCTGCCGGGAGTAAAGGCCCTCGTCTAT GTCTGCTTCACTGCCGTTCTTGGCCACTCCCTAGCAATAGAGGAAAAGAACAAGTTTAGGG*A*G-3′ |
| | VS (M41T) | 5′-T*C*CACACCCCTCAGCCCGGCCTTGGCCCCACTTACAGCTGCCGGGAGTAAAGGCCCTCGTCTAT GTCTGCTTCACTGCCGTTCTTGGCCGTTCCCTAGCAATAGAGGAAAAGAACAAGTTTAGGG*A*G-3′ |
| | VS (M41L) | 5′-T*C*CACACCCCTCAGCCCGGCCTTGGCCCCACTTACAGCTGCCGGGAGTAAAGGCCCTCGTCTAT GTCTGCTTCACTGCCGTTCTTGGCCAGTCCCTAGCAATAGAGGAAAAGAACAAGTTTAGGG*A*G-3′ |
| | Unmutated (M41) | 5′ -T*C*CACACCCCTCAGCCCGGCCTTGGCCCCACTTACAGCTGCCGGGAGTAAAGGCCCTCGTCTAT GTCTGCTTCACTGCCGTTCTTGGCCATCCCCTAGCAATAGAGGAAAAGAACAAGTTTAGGG*A*G-3′ |
| sgRNA | | 5′ -GCCGTTCTTGGCCATTCCCT-3′ |

sgRNAs were designed to target the *UBA1* gene sequence surrounding the M41 codon. *Phosphorothioate backbone modification (PS)

**Supplemental Table 10. Antibodies used in immunoblotting.**

| Type | Antibody | Cat. No. | Source |
|---|---|---|---|
| Primary | UBE1L2/UBA6 Antibody | 13386 | Cell Signaling Technologies |
| | Ubiquitin (P4D1) Mouse mAb | 3936S | Cell Signaling Technologies |
| | UBE1a/b Antibody | 4891 | Cell Signaling Technologies |
| | Anti-GAPDH antibody (6C5) | ab8245 | Abcam |
| | Anti-Lamin B1 Antibody | ab16048 | Abcam |
| | Phospho-MLKL (Ser358) (D6H3V) Rabbit mAb | 91689S | Cell Signaling Technologies |
| | Anti-MLKL antibody (EPR17514) | ab184718 | Abcam |
| | Anti-RIP3 (phospho S227) antibody (EPR9627) | ab209384 | Abcam |
| | RIP3 (E7A7F) XP® Rabbit mAb | 10188 | Cell Signaling Technology |
| | Phospho-RIPK1 (Ser166) Polyclonal antibody | 28252-1-AP | Proteintech |
| | RIPK1 Polyclonal antibody | 29932-1-AP | Proteintech |
| Secondary | Goat anti-Mouse IgG (H+L) Secondary Antibody, HRP | 31430 | Invitrogen |
| | Goat anti-Rabbit IgG (H+L) Secondary Antibody, HRP | 31460 | Invitrogen |

**Supplemental Table 11. Primers used in PCR, Sanger sequencing, and RT-qPCR.**

| | Target gene | Forward | Reverse |
|---|---|---|---|
| Genotyping primer | UBA1 (WT) | AGGTCTGACAGCTGTGTTACCCTGG | GCCGTTCTTGGCCATTCCCTAGGAA |
| | *UBA1* (M41V) | AGGTCTGACAGCTGTGTTACCCTGG | CCGTTCTTGGCCACTCCCTAGC |
| | *UBA1* (M41T) | AGGTCTGACAGCTGTGTTACCCTGG | CCGTTCTTGGCCATCCCCTAGC |
| | *UBA1* (M41L) | AGGTCTGACAGCTGTGTTACCCTGG | CCGTTCTTGGCCAGTCCCTAGC |
| | *UBA1* (+ silent mutation for unmutated cells) | AGGTCTGACAGCTGTGTTACCCTGG | CCGTTCTTGGCCGTTCCCTAGC |
| Sequence primer | *UBA1* (Met41 amplification from genomic DNA) | CGGTACCCATGTGCTCCAGGGTC | GGTAACAGCCTTGACCCCACCAAGG |
| | *UBA1* (Met41 amplicon sequencing) | CCGCTGTCCAAGAAACGTC | |
| qPCR Primer | *UBA1B* (exogenous) | CGATACACCATCCGCTGAACGCGTG | AGAGGCCACTTGTGTAGCGCCAAG |
| | *GAPDH* | GAAATCCCATCACCATCTTCCAGG | GAGCCCCAGCCTTCTCCATG |
| | *XBP1u* | GGAGTTAAGACAGCGCTTGGGGATG | TGTTCTGGAGGGGTGACAACTGGG |
| | *XBP1s* | GCTGAGTCCGCAGCAGGT | GGCTCTGGGGAAGGGCATTT |
| | *IL1B* | TGCACGATGCACCTGTACGA | GAGAACACCACTTGTTGCTCCA |
| | *RNASE1* | GCTTTTCTGGGAAAGTGAGGCCAC | CCAGCACCAGCAGTATCAGGACAAG |
| | *IFI27* | CCTCCATAGCAGCCAAGATGATGTC | GGATGAACTTGGTCAATCCGGAGA |
| | *MX1* | GTTGGAGGCACTGTCAGGAGTTG | GCCTCTCCACTTATCTTCGTTCACA |
| | *CXCL10* | GCAAGCCAATTTTGTCCACGTGTTGAGATC | ACTGCATCGATTTTGCTCCCCTCTGG |

**Supplemental Table 12. shRNA sequences used for hRNASE1 knockdown**.

| Name | Sequence |
|---|---|
| shRNA#1 | 5' -TCCTTCTGCTTGTCCTGATACCTCGAGGTATCAGGACAAGCAGAAGGA -3' |
| shRNA#2 | 5' -GAAATTCCAGCGGCAGCATATCTCGAGATATGCTGCCGCTGGAATTTC -3' |
| shRNA#3 | 5' -TGCAAACCAGTGAACACCTTTCTCGAGAAAGGTGTTCACTGGTTTGCA -3' |
| shRNA-control (scrambled) | 5' -CCTAAGGTTAAGTCGCCCTCGCTCGAGCGAGGGCGACTTAACCTTAGG -3' |

### Supplemental Figures

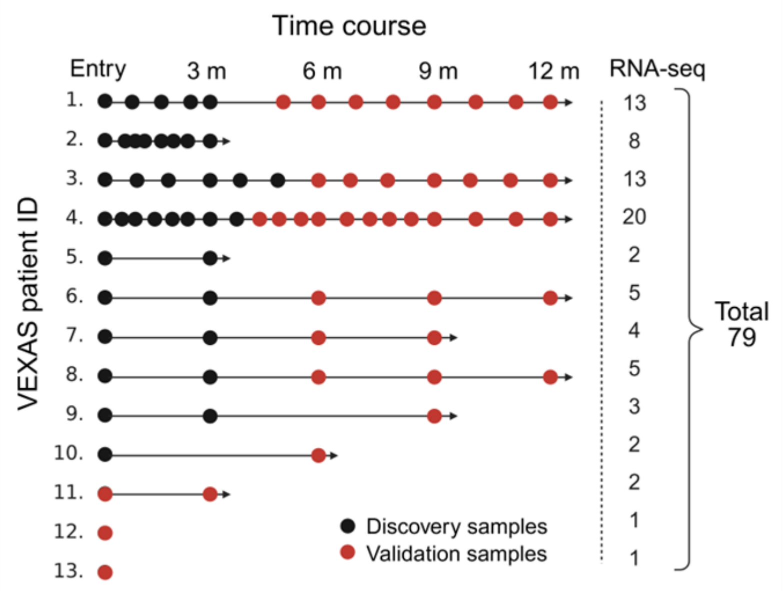


**Supplemental Figure 1. Timeline of sampling during the deep phenotyping study.**

Peripheral blood sampling times for 13 patients with VS (ID: 1–13) are shown, divided into discovery and validation phases. Numbers on the right indicate the total RNA-seq sample count for each patient.

VS, vacuole, E1 enzyme, X-linked, autoinflammatory, somatic (VEXAS) syndrome.

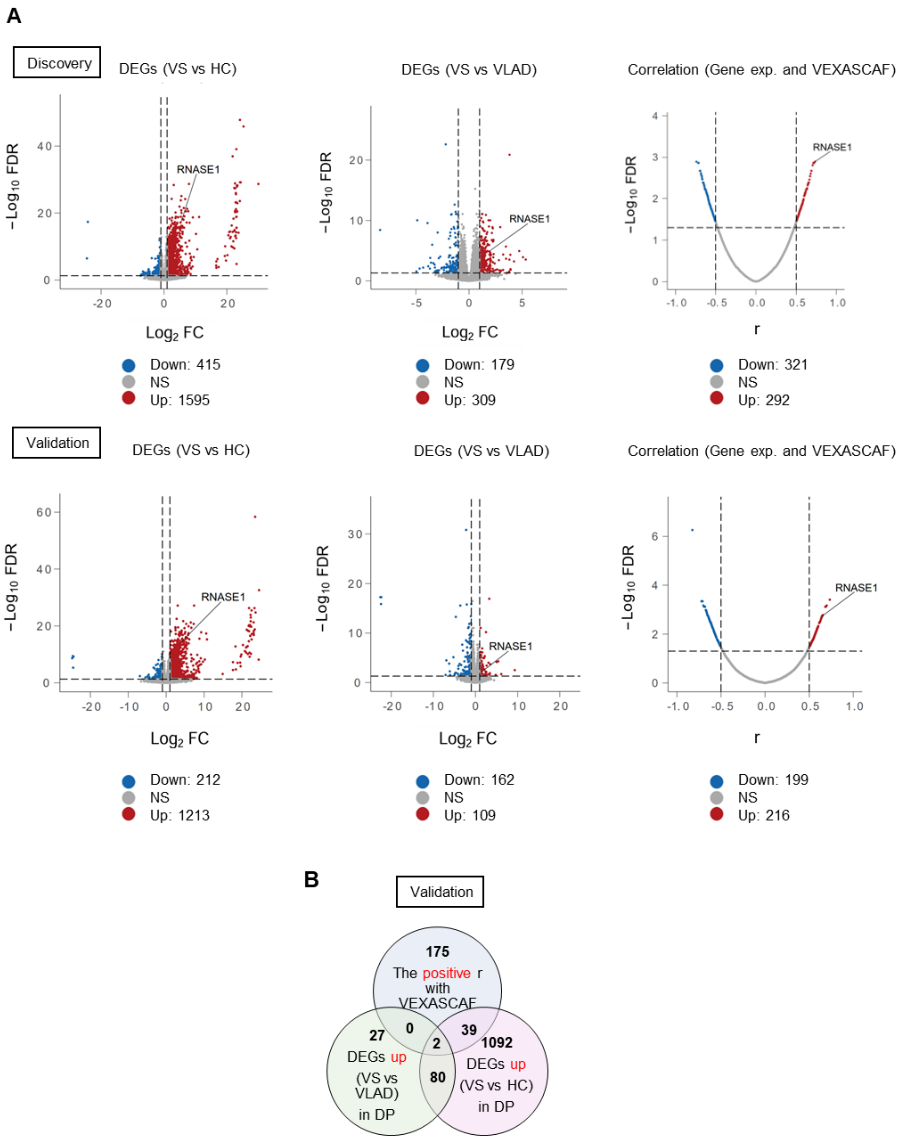


**Supplemental Figure 2. The process of identifying RNASE1 through DP.**

**A:** Volcano plots of DEGs in "VS vs VLAD," "VS vs HC," and genes correlated with VEXASCAF from the discovery and validation cohorts in DP.

**B:** Venn diagram of the validation cohort. *RNASE1* exhibits the highest correlation coefficient among the two genes.

DEGs were defined as FC > 2, FC < 1/2, and FDR < 0.05. DEGs, differentially expressed genes; DP, deep phenotyping; HC, healthy controls; VEXASCAF, VEXAS Current Activity Form; VLAD, VS-like autoinflammatory diseases; VS, vacuole, E1 enzyme, X-linked, autoinflammatory, somatic (VEXAS) syndrome.

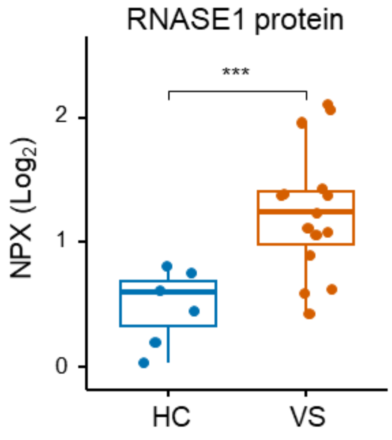


**Supplemental Figure 3. Elevated RNASE1 expression in patients with VS compared with HC determined using plasma proteome analysis.**

Plasma RNASE1 protein levels were assessed in patients with VS and HC using high-sensitivity proteomic analysis (Olink). (HC, n = 3; VS, n = 5; technical duplicates or triplicates were included).

*$P < 0.05$, **$P < 0.01$, ***$P < 0.001$. Box-and-whisker plots represent the median, interquartile range (IQR), and 1.5× IQR. Statistical comparisons were performed using the two-sided Mann–Whitney U test.

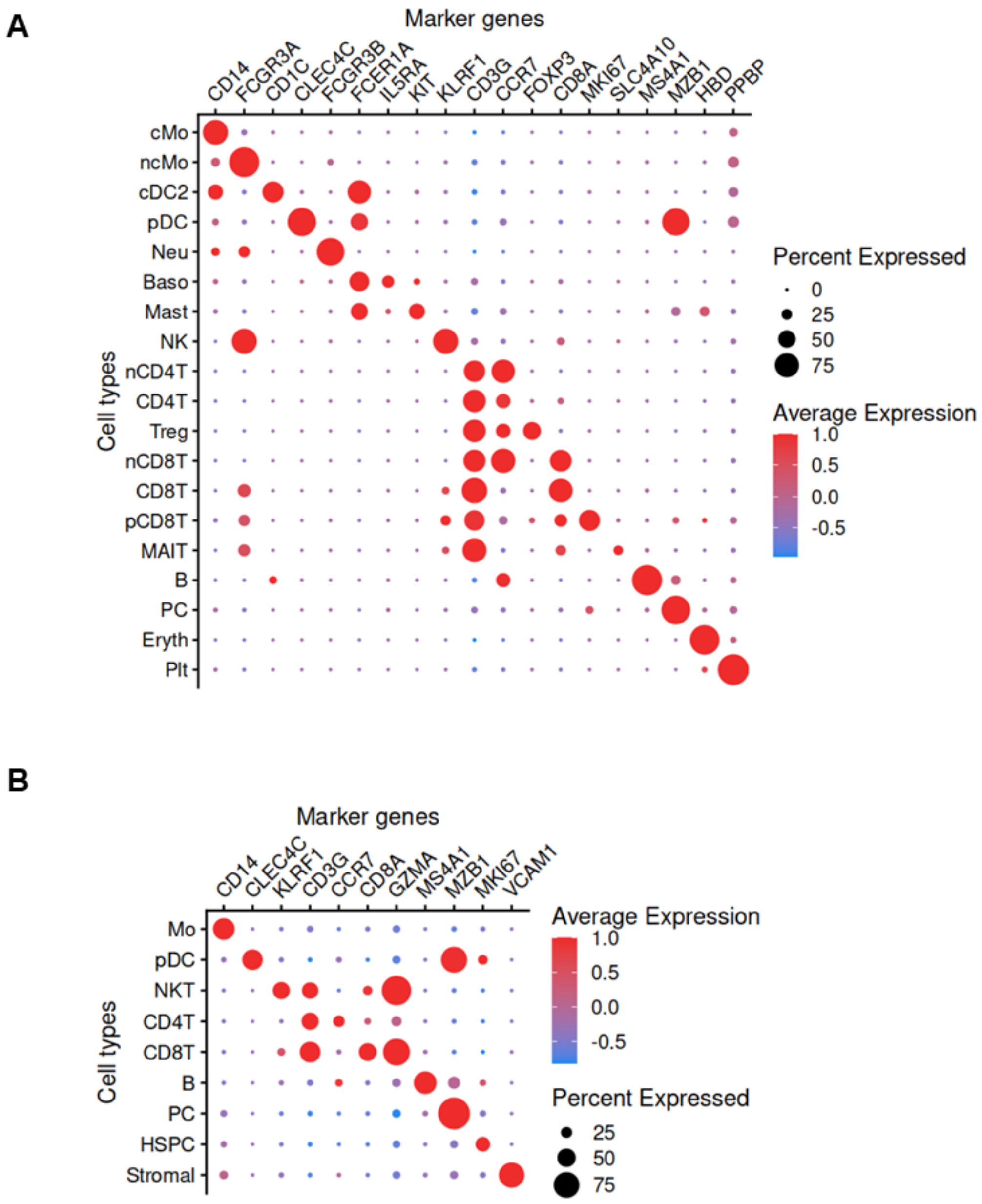


**Supplemental Figure 4. Single-cell RNA sequencing cluster annotations.**

**A, B:** Bubble plot showing representative marker genes used to identify and annotate each cell-type cluster in the scRNA-seq datasets. (**A**) PBMCs from YCU (HC; n = 1 and VS; n = 2) and a public dataset (Mizumaki et al., HC; n = 5 and VS; n = 9) and (**B**) bone marrow mononuclear cells from a public dataset (Wu et al., HC; n = 4 and VS; n = 9).

Bubble size indicates the percentage of cells expressing the gene, and color intensity represents the average expression level.

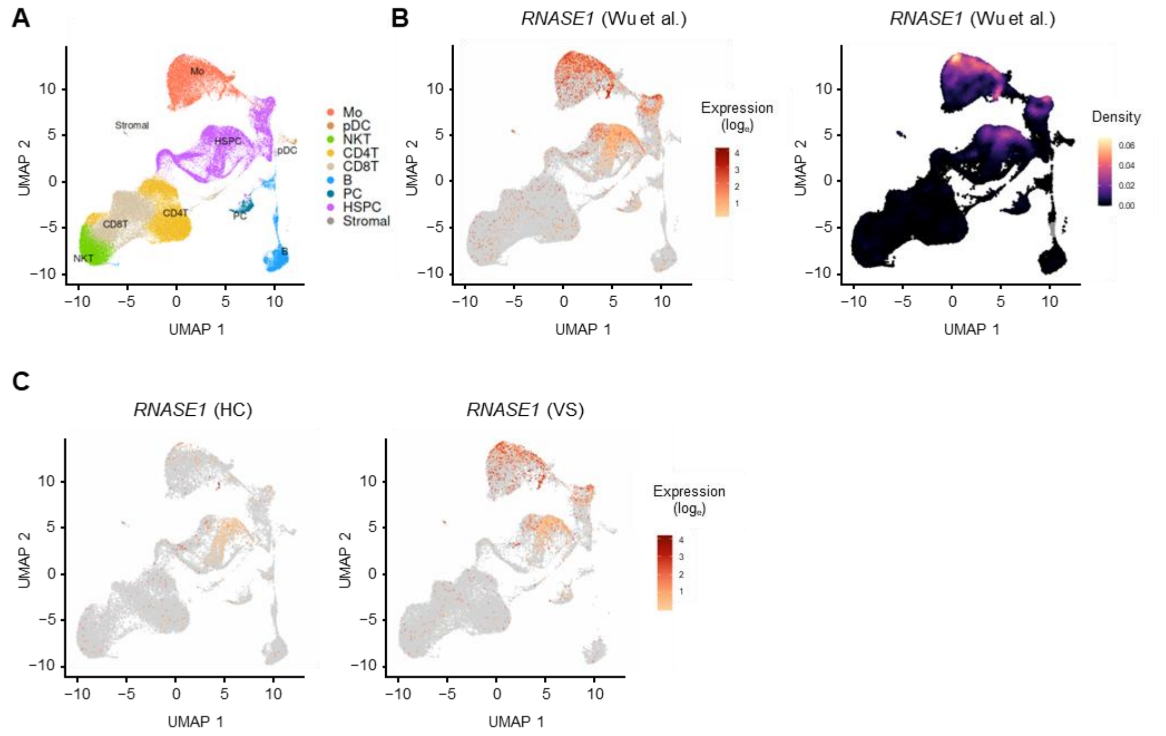


**Supplemental Figure 5. Single-cell RNA sequencing of bone marrow mononuclear cells.**

**A-B:** UMAP plot of 121,864 bone marrow mononuclear cells (BMMCs) from all subjects (n = 13; Wu et al.). (**A**) Cell type annotations for each cluster. Mo: monocyte, pDC: plasmacytoid dendritic cell, NKT: natural killer T, PC: plasma cell, HSPC: hematopoietic stem and progenitor cell. (**B**) UMAP by *RNASE1* expression levels (left) and cell density (right).

**C:** UMAP plots of BMMCs derived from HC (n = 4) and VS patients (n = 9), colored according to *RNASE1* expression levels in each dataset. The total cell numbers were downsampled to 30,000 cells per group.

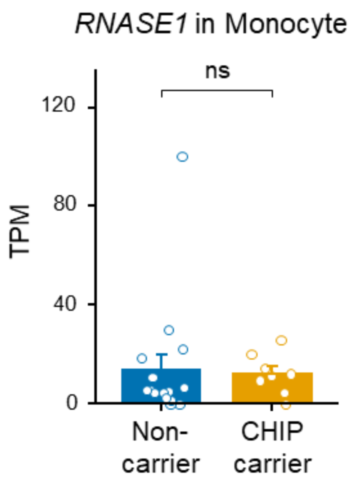


**Supplemental Figure 6. Comparison of *RNASE1* expression in elderly peripheral monocytes with or without clonal hematopoiesis.**

In a public monocyte RNA-seq dataset (Dederichs TS et al.) of older adult patients undergoing heart surgery or carotid endarterectomy, *RNASE1* expression did not differ between clonal hematopoiesis of indeterminate potential (CHIP) carriers (n = 8) and non-carriers (n = 16).

ns: not significant. Error bars indicate the standard error of the mean. Statistical comparisons were performed using the two-sided Mann–Whitney U test.

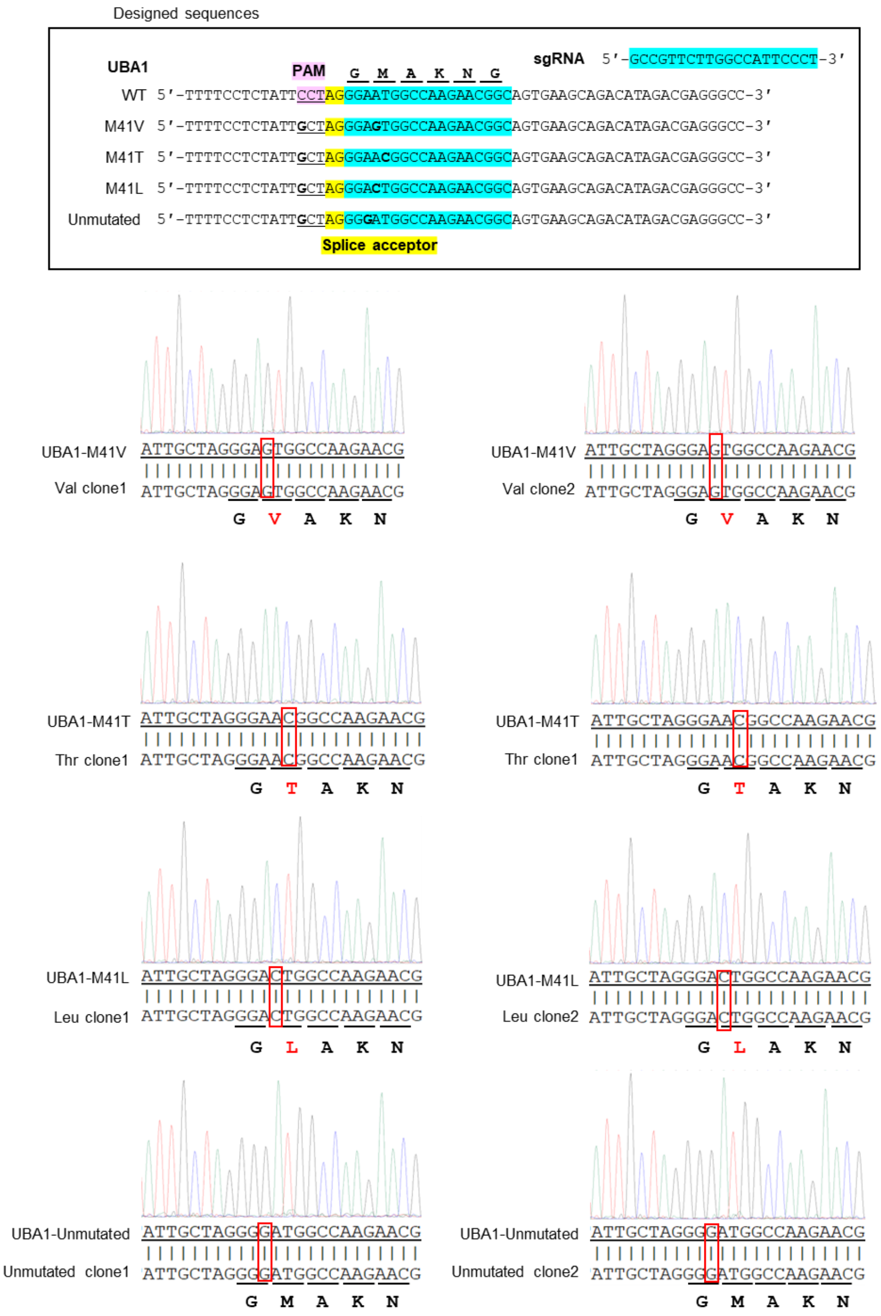


**Supplemental Figure 7. Sanger sequencing of PCR products from genomic DNA isolated from VS and unmutated cell lines.**

*UBA1* genotypes of the single-cell clones were confirmed by Sanger sequencing. These mutations are highlighted, including the M41 mutations in *UBA1* (M41V, M41T, and M41L), as well as a mutation introduced in the PAM sequence to disrupt its function.

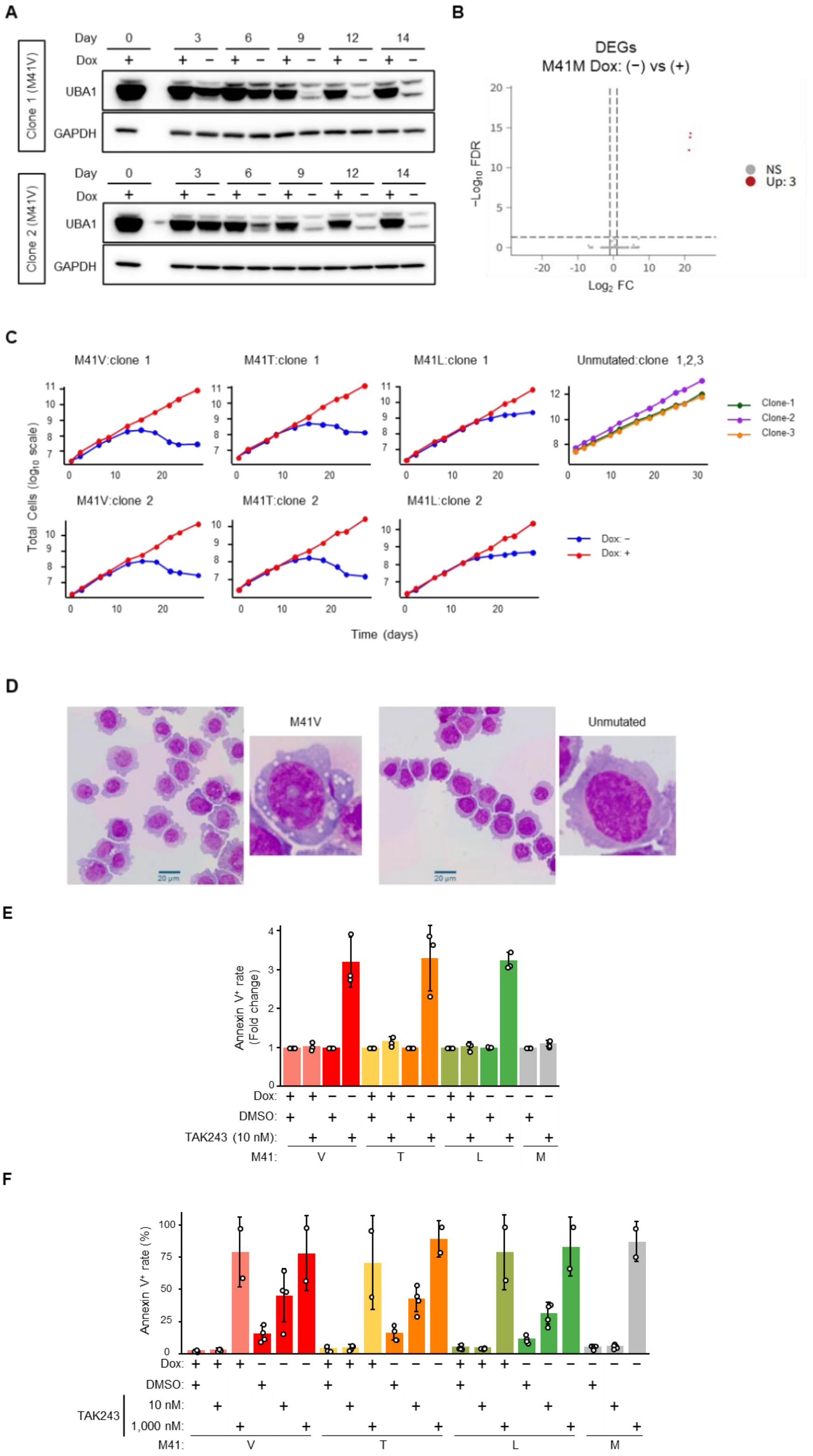

A
Day
0
3
6
9
12
14
Dox
Clone 1 (M41V)
UBA1
GAPDH
Clone 2 (M41V)
B
DEGs
M41M Dox: (−) vs (+)
−Log10 FDR
Log2 FC
NS
Up: 3
C
M41V:clone 1
M41T:clone 1
M41L:clone 1
Unmutated:clone 1,2,3
Clone-1
Clone-2
Clone-3
M41V:clone 2
M41T:clone 2
M41L:clone 2
Dox: −
Dox: +
Total Cells (log10 scale)
Time (days)
D
M41V
Unmutated
20 µm
E
Annexin V+ rate (Fold change)
Dox:
DMSO:
TAK243 (10 nM):
M41:
V
T
L
M
F
Annexin V+ rate (%)
Dox:
DMSO:
TAK243
10 nM:
1,000 nM:
M41:
V
T
L
M

**Supplemental Figure 8. Phenotypic characterization of VS cell lines.**

**A:** Immunoblot analysis of UBA1 isoforms in two indicated M41V cell clones during differentiation from day 0 to day 14, cultured in medium containing 20 ng/mL doxycycline (Dox) (Tet-On) or in Dox-free medium (Tet-Off).

**B:** Volcano plots of DEGs from RNA-seq of unmutated (M41M) cell lines under Tet-Off and Tet-On conditions. Three clones with four biological replicates (n = 12) for Tet-Off, and three clones with two biological replicates (n = 6) for Tet-On, were analyzed. The experiments were conducted 18 days after switching. DEGs were defined as FC > 2, FC < 1/2, and FDR < 0.05.

DEGs, differentially expressed genes; FC, fold change; FDR, false discovery rate.

**C:** Growth curves of cell lines harboring *UBA1* mutations (M41V, M41T, or M41L). VS cell lines cultured under Tet-On conditions are shown in red, whereas those cultured under Tet-Off conditions are shown in blue. The continued proliferation of VS cell lines for approximately two weeks after switching to Tet-Off is likely due to the time required for the degradation of exogenous UBA1b protein. All unmutated cell lines were cultured under Tet-Off conditions.

**D:** Representative images of M41V and unmutated cell lines stained with Wright–Giemsa 13 days after switching.

**E, F:** Bar plots show the extent of Annexin V-positive cell death in VS and unmutated cell lines treated with DMSO or TAK-243 (10 nM, 1,000 nM). At low concentrations, TAK-243 exhibited high sensitivity in the VS cell lines; however, at high concentrations, it triggered cell death in Tet-On cells. The experiments were conducted on day 19 after switching to Tet-Off condition. Error bars indicate the standard deviation (SD). Data represent a compilation of 2–3 independent experiments using different clones (**E** and **F**).

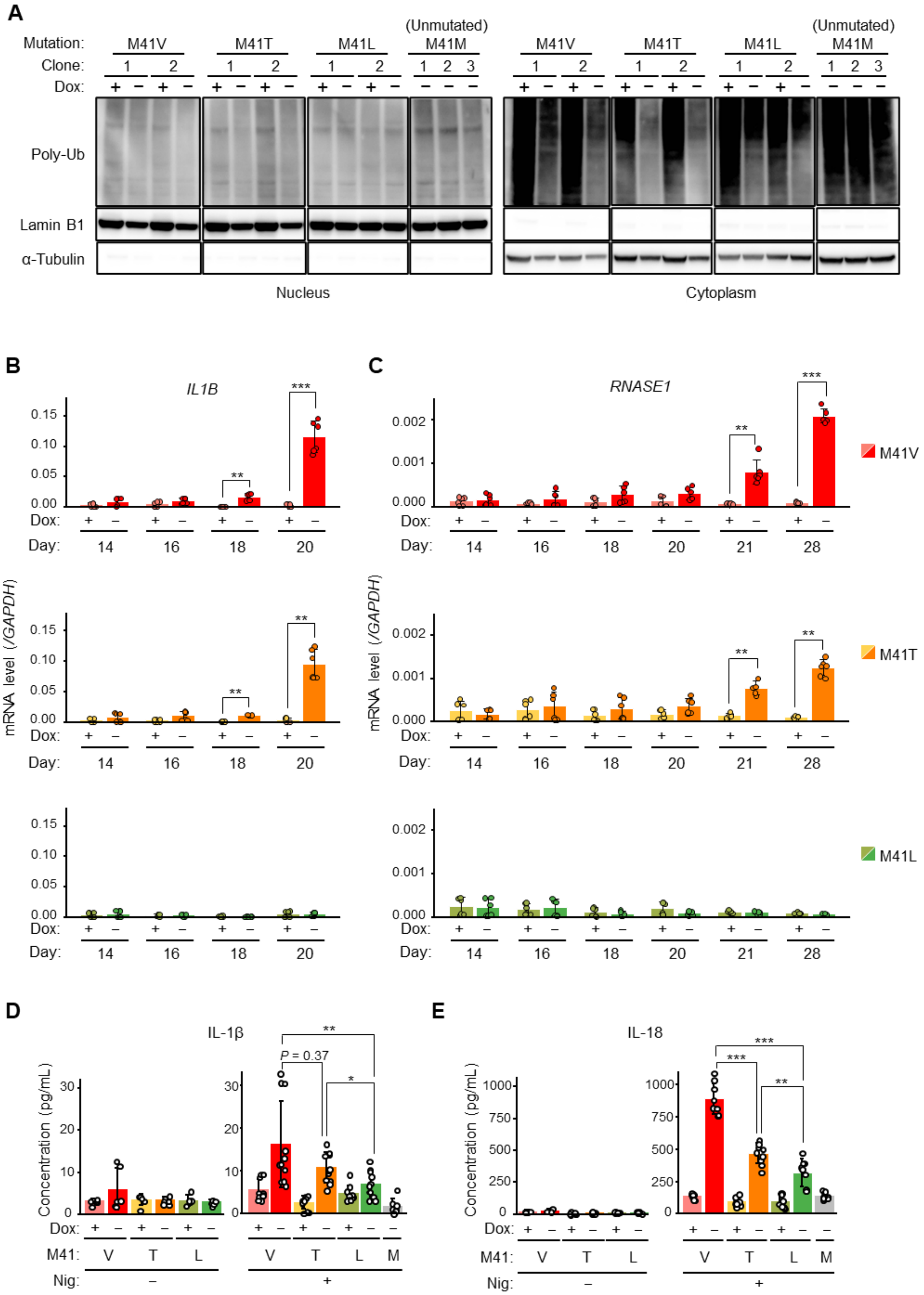
A
Mutation:
M41V
M41T
M41L
(Unmutated) M41M
Clone:
Dox:
Poly-Ub
Lamin B1
α-Tubulin
Nucleus
Cytoplasm
B
IL1B
C
RNASE1
M41V
M41T
M41L
mRNA level (/GAPDH)
Dox:
Day:
D
IL-1β
P = 0.37
Concentration (pg/mL)
M41:
Nig:
E
IL-18

**Supplemental Figure 9. VS cell lines exhibit genotype-dependent phenotypic differences.**

**A:** Immunoblot analysis of polyubiquitin proteins after nuclear and cytoplasmic fractionation. Lamin B1 (nuclear marker) and α-tubulin (cytoplasmic marker) were used as loading controls. The experiments were conducted on day 19 after switching to Tet-Off.

**B, C:** mRNA expression levels of *IL1B* (**B**) and *RNASE1* (**C**) in VS cell lines under Tet-On/Off conditions measured on different culture days.

**D, E:** Release of IL-1β (**D**) and IL-18 (**E**) into the supernatant from VS cell lines after stimulation with 10 μM nigericin (Nig) for 2 h (right). Without nigericin stimulation, neither IL-1β nor IL-18 was detected (left panel). The experiments were conducted 15 days after switching.

$^{*}P < 0.05$, $^{**}P < 0.01$, $^{***}P < 0.001$, ns: not significant. Data are shown as means ± SD. *P*-values were determined using a two-sided t-test (**B-E**). Data are representative of three independent repeat experiments (**A**). Data were pooled from 2–4 independent experiments using different clones (**B-E**). IL, interleukin.

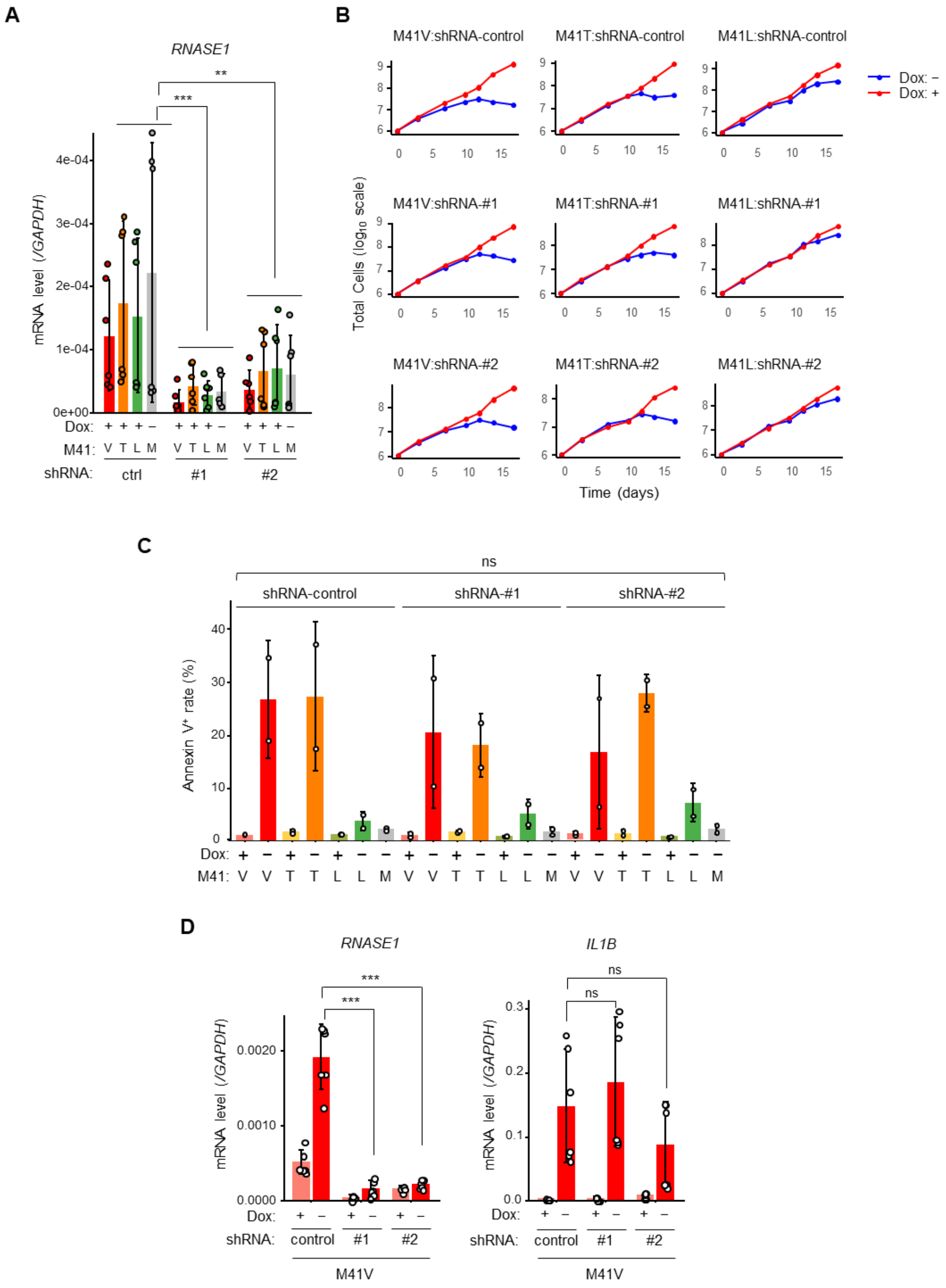


**Supplemental Figure 10. Effect of RNASE1 knockdown on cell death and inflammatory gene expression in VS cell lines.**

**A:** qPCR analysis of *RNASE1* mRNA expression to confirm RNASE1 knockdown. shRNA#3 was excluded from the experiments because of its low knockdown efficiency. The experiments were performed using Tet-On cells expressing *UBA1* mutations (M41V, M41T, and M41L), as well as Tet-Off unmutated cells.

**B:** Growth curves of cell lines harboring *UBA1* mutations (M41V, M41T, and M41L) transduced with *RNASE1* shRNA. VS cell lines cultured in a medium containing Dox (Tet-On) are shown in red, whereas those cultured in a Dox-free medium (Tet-Off) are shown in blue. Compared to Supplemental Figure 8C, cell death under Tet-Off conditions occurred earlier, likely because of the lower Dox concentration used during cell stock preparation. All unmutated cell lines were maintained under Tet-Off conditions.

**C:** Annexin V-positive rates of VS and unmutated cell lines transduced with *RNASE1* shRNA, assessed on day 18 post-switching, shown as bar plots.

**D:** *RNASE1* and *IL1B* mRNA expression levels in M41V cell lines transduced with *RNASE1* shRNA. Although *RNASE1* expression was significantly reduced, *IL1B* expression remained unaltered. The experiments were conducted 18 days after switching.

$^{*}P < 0.05$, $^{**}P < 0.01$, $^{***}P < 0.001$, ns: not significant. The mean and SD are shown (**A, C, D**). *P*-values were determined using a two-sided t-test (**A**, **D**) and the two-sided Kruskal-Wallis test for multiple groups (#1, #2, ctrl) comparisons (**C**). Data were pooled from two independent experiments using different clones (**A, C, D**).

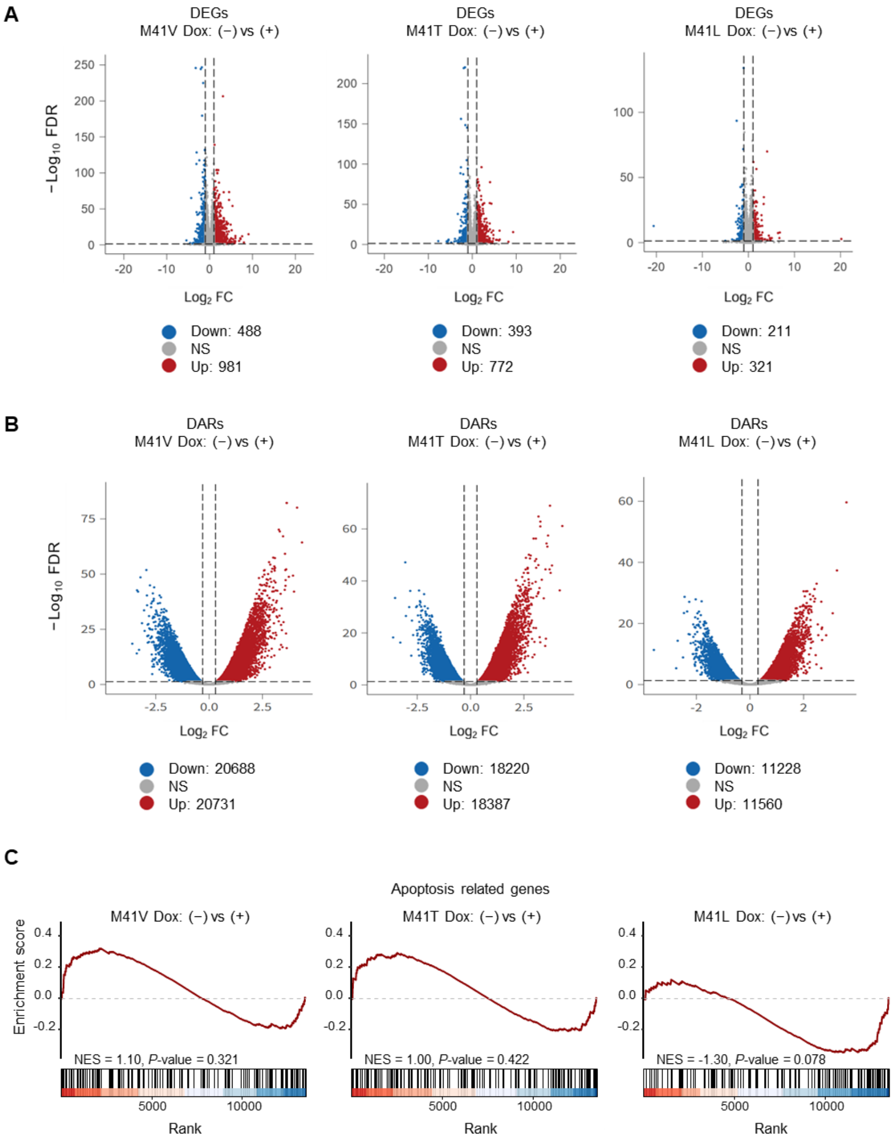


**Supplemental Figure 11. Multi-omics analysis of VS cell lines.**

**A:** Volcano plots of DEGs in each genotype from RNA-seq of VS cell lines under Tet-Off vs. Tet-On conditions. DEGs were defined as FC > 2, FC < 1/2, and FDR < 0.05.

**B:** Volcano plots of DARs in each genotype from ATAC-seq of VS cell lines under Tet-Off vs. Tet-On conditions. DARs were defined as FC > 1.5, FC < 1/1.5, and FDR < 0.05. DARs, differentially accessible regions.

**C:** Gene Set Enrichment Analysis of the apoptosis-related hallmark gene set (HALLMARK_APOPTOSIS), comparing Tet-Off vs. Tet-On in VS cell lines with different *UBA1* genotypes.

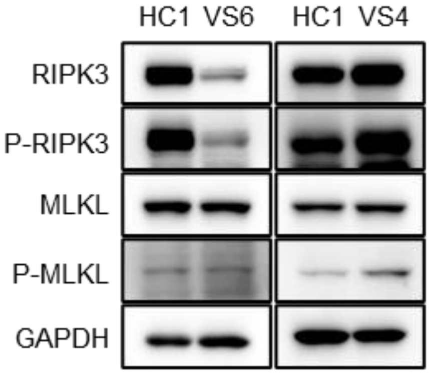


**Supplemental Figure 12. MLKL phosphorylation in primary VS monocytes.**

Primary monocytes from a healthy control (HC1) and patients with VS (IDs: 6 and 4) were subjected to immunoblotting for necroptosis associated proteins. Elevated p-MLKL was detected in the high-activity patient (ID: 4, VEXASCAF 2), but not in the low-activity patient (ID: 6, VEXASCAF 0).

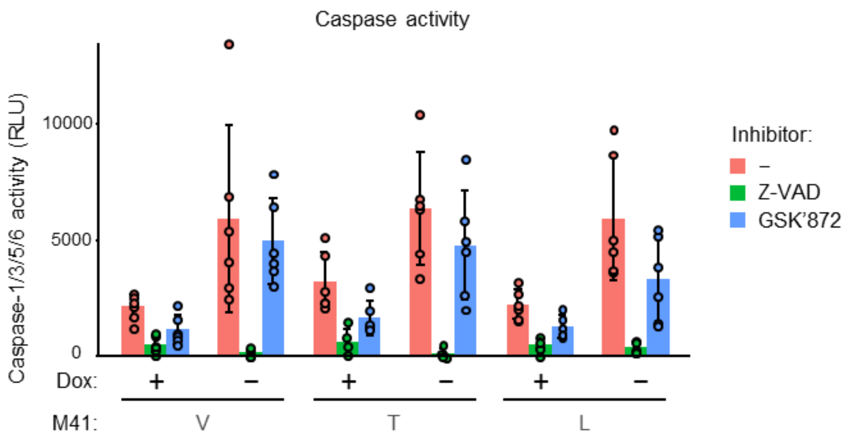


**Supplemental Figure 13. Caspase activity assay of VS cell lines treated with cell death pathway inhibitors.**

Caspase activity was measured in the VS cell lines cultured with the respective inhibitors. Treatment with Z-VAD markedly suppressed the activity of Caspases-1, -3, -5, and -6. The experiments were conducted 22 days after switching.

The mean and SD are shown. Data were pooled from three independent experiments.

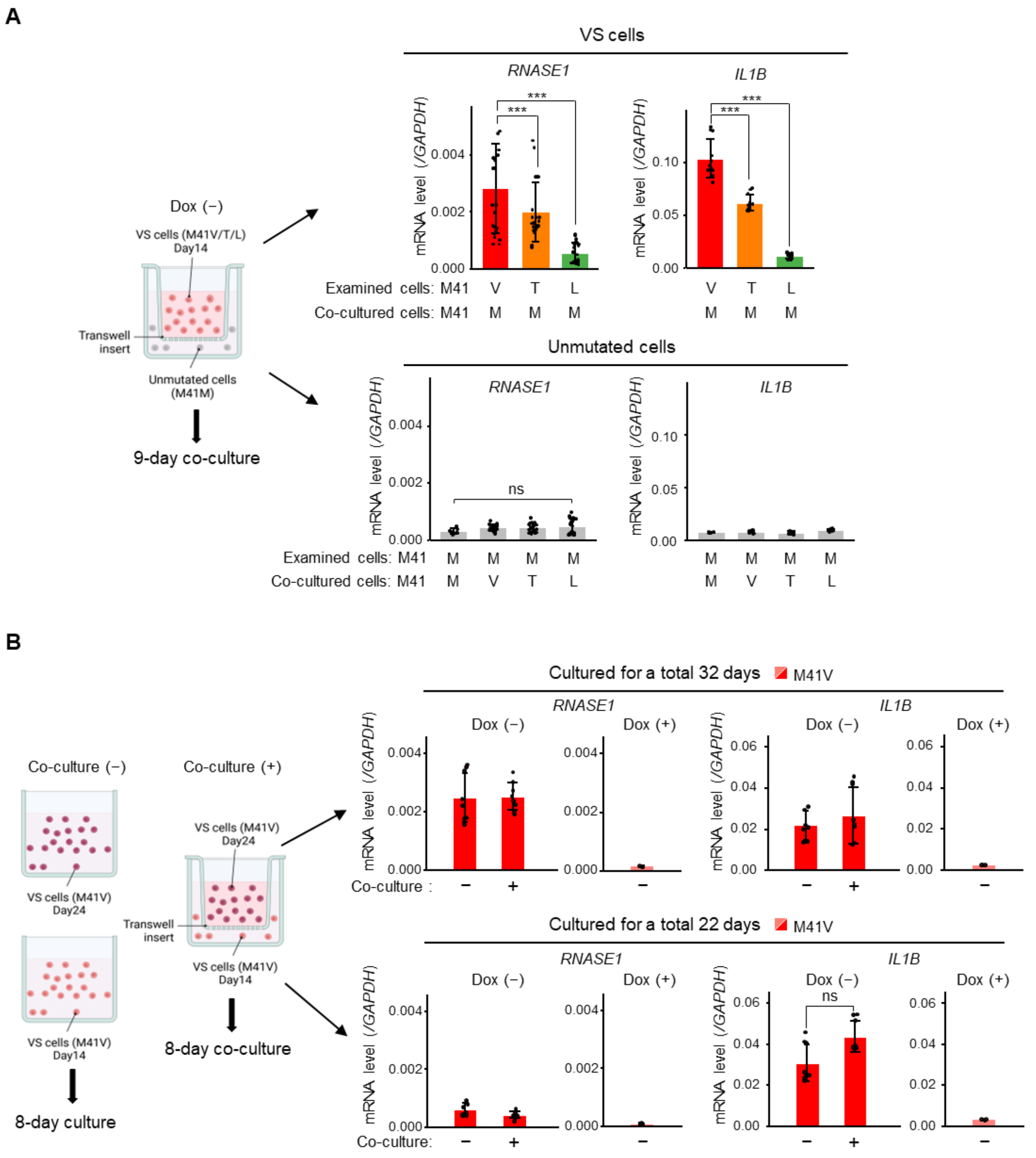


**Supplemental Figure 14. Effect of damage-associated molecular patterns on *RNASE1* and *IL1B* expressions assessed by transwell assay.**

**A:** Schematic of transwell assay (left) and targeted gene expression analysis of cells cultured under Tet-Off conditions (right). VS (M41V, M41T, M41L) cell lines, pre-cultured for 14 days after Dox deprivation were seeded in the upper well, and unmutated (M41M) cell lines were seeded in the lower well. The cells were co-cultured for 9 days.

**B:** Schematic of the transwell assay (left) and results of targeted gene expression analysis (right). VS (M41V) cell lines, pre-cultured for two different durations (14 and 24 days) after Dox deprivation, were co-cultured for 8 days, resulting in total culture durations of 22 and 32 days, respectively. *RNASE1* and *IL1B* expression levels in co-cultured cells were compared with those in cells cultured alone under the same total culture duration.

$^{*}P < 0.05$, $^{**}P < 0.01$, $^{***}P < 0.001$, ns: not significant. The mean and SD are shown. *P*-values were determined using a two-sided t-test. Data were pooled from two independent experiments using different clones.
